# Estrogenic control of reward prediction errors and reinforcement learning

**DOI:** 10.1101/2023.12.09.570945

**Authors:** Carla E. M. Golden, Audrey C. Martin, Daljit Kaur, Andrew Mah, Diana H. Levy, Takashi Yamaguchi, Amy W. Lasek, Dayu Lin, Chiye Aoki, Christine M. Constantinople

**Affiliations:** Center for Neural Science, New York University; New York, NY 10003.; Neuroscience Institute, New York University Grossman School of Medicine; New York, NY 10016.; Department of Pharmacology and Toxicology, Virginia Commonwealth University; Richmond, VA 23298.

## Abstract

Gonadal hormones act throughout the brain^1^, and neuropsychiatric disorders vary in symptom severity over the reproductive cycle, pregnancy, and perimenopause^2–4^. Yet how hormones influence cognitive processes is unclear. Exogenous 17*β*-estradiol modulates dopamine signaling in the nucleus accumbens core (NAcc)^5,6^, which instantiates reward prediction errors (RPEs) for reinforcement learning^7–16^. Here we show that endogenous 17*β*-estradiol enhances RPEs and sensitivity to previous rewards by reducing dopamine reuptake proteins in the NAcc. Rats performed a task with different reward states; they adjusted how quickly they initiated trials across states, balancing effort against expected rewards. NAcc dopamine reflected RPEs that predicted and causally influenced initiation times. Elevated endogenous 17*β*-estradiol increased sensitivity to reward states by enhancing dopaminergic RPEs in the NAcc. Proteomics revealed reduced dopamine transporter expression. Finally, knockdown of midbrain estrogen receptors suppressed reinforcement learning. 17*β*-estradiol therefore controls RPEs via dopamine reuptake, mechanistically revealing how hormones influence neural dynamics for motivation and learning.

## Introduction

Estrogenic hormones, including 17*β*-estradiol, rhythmically fluctuate over the reproductive cycle in mammalian females and act on estrogen receptors, which can regulate protein expression via transcription and activation of signal transduction pathways^1,17,18^. These hormonal fluctuations support physiological imperatives such as homeostasis, fertility, and sexual behavior^1^, but due to broad and diffuse mechanisms of action, also bind to receptors in brain regions that support cognitive behaviors, including decision-making.

One important neural system modulated by estrogen receptors is dopamine signaling in the nucleus accumbens core (NAcc), which is modulated by application of exogenous 17*β*-estradiol in slices of ovariectomized animals^5,19^. The NAcc is a part of the ventral striatum that is innervated by midbrain dopaminergic neurons in the ventral tegmental area (VTA), and is required for learning cue-outcome associations^20^ and determining the vigor of motivated behaviors^21^. Dopamine release in the NAcc is thought to instantiate RPEs for reinforcement learning, which is a powerful theoretical framework for describing how animals learn through interacting with the environment^22^.

In reinforcement learning, agents learn the value of taking actions in different states; those state-action values are iteratively updated based on RPEs, or the difference between received and expected rewards. A wealth of evidence suggests that dopamine release in the NAcc encodes RPEs^7–12^. Critically, manipulating dopamine release in this circuit has provided causal evidence that it acts as an RPE for reinforcement learning^13–16^. More precisely, converging synaptic inputs (e.g., from cortex, hippocampus) onto medium spiny neurons, the principal cells of the striatum, are thought to convey the state the animal is in. Dopamine is thought to mediate plasticity at these synapses, such that animals will be more (or less) likely to take actions in a state that produced positive (or negative) RPEs. In actor-critic models of the basal ganglia, the NAcc is thought to act as a “critic” that learns the expected future reward in a given state (i.e., state value)^23,24^, which is known to influence response vigor^25–28^. Therefore, RPEs in the NAcc, in addition to training animals to take actions that will maximize future rewards, likely influence whether actions in a given state will be more or less vigorous^21^. The fact that 17*β*-estradiol can influence dopamine release in the NAcc suggests that hormones may influence RPEs for reinforcement learning. However, how behaviors unrelated to reproduction that are critical to survival, such as reward-driven learning, respond to endogenous fluctuations in 17*β*-estradiol is unresolved.

Here we find that rats update their response vigor on a trial-by-trial basis based on dopamine RPEs in the NAcc. We show that estrogen receptors in the midbrain enhance reinforcement learning in the stage of the reproductive cycle when 17*β*-estradiol is high by suppressing expression of dopamine transporters in the NAcc, which increases the dynamic range of dopaminergic RPEs. This reveals a novel role for dopamine reuptake mechanisms in shaping RPEs for reinforcement learning.

## Results

### 17*β*-estradiol enhances learning

We trained rats on a self-paced temporal wagering task that manipulated reward expectations by varying the magnitude of offered rewards in blocks of trials^28^ (Fig. 1a,b). Rats initiated trials by poking their nose in a center port. This triggered an auditory cue, the frequency of which indicated the volume of the water reward offered on that trial (4, 8, 16, 32, or 64 *µ*L). The reward was assigned randomly to one of two side ports, indicated by an LED. The rat could wait for an uncued and unpredictable delay to obtain the reward, or at any time could opt out by poking in the other side port to start a new trial. On 15-25% of trials, rewards were withheld to assess how long rats were willing to wait for them (catch trials).

**Figure 1:**
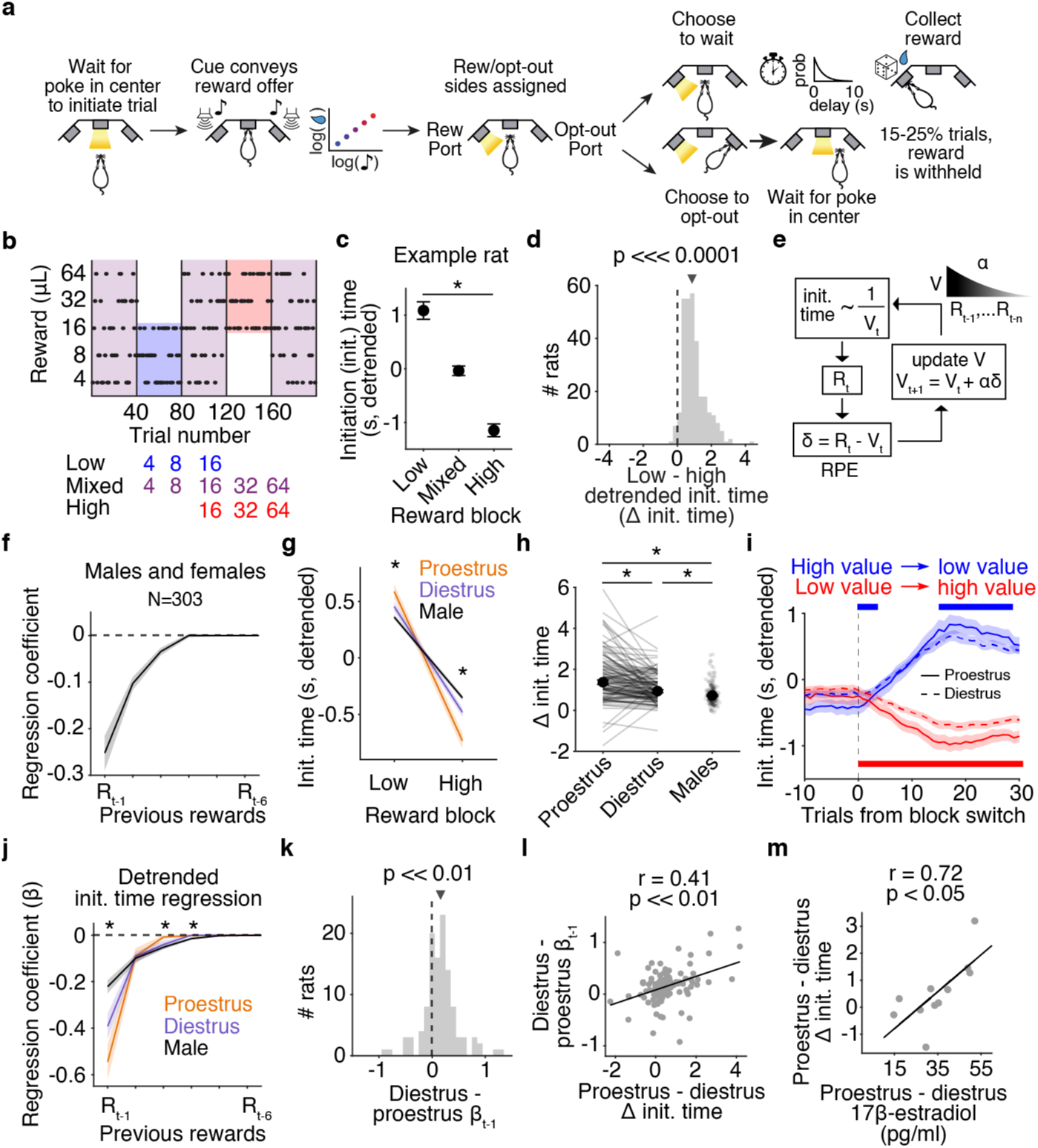
Rats’ trial initiation times are modulated by the value of the state and estrous stage. **a.** Schematic of behavioral paradigm. **b.** Block structure of task with an example session (top) and reward distributions in each block (bottom). **c.** Mean detrended trial initiation times across blocks for one example rat. Initiation times in low and high blocks are significantly different, *p <<* 1 × 10^−20^, two-sided Wilcoxon rank sum test, error bars are confidence intervals (CIs). **d.** Sensitivity of detrended initiation times to blocks (low - high block) across the population is significantly different from zero, one-sided Wilcoxon signed rank test *p <<* 1 × 10^−20^, *N* = 303. **e.** Schematic depicting reinforcement learning model. Initiation times are inversely proportional to state value (*V_t_*) on trial (*t*), which gets updated by a reward prediction error (*δ*) when a reward is offered on that trial (*R_t_*). A learning rate (*α*) determines how much to weigh the previous rewards (*R*_t-n_) in the estimation of state values. **f.** Median regression coefficients of detrended trial initiation times as a function of rewards on previous trials during mixed blocks for all rats. **g.** Median detrended initiation times for low and high blocks, *N* = 118 female rats in proestrus and diestrus and *N* = 185 male rats (Kruskal–Wallis test *p* = 1.04 × 10^−6^ for low and *p* = 3.96×10^−14^ for high blocks, two-sided Wilcoxon signed-rank test *p* = 1.38×10^−5^ and *d* = 0.38 for low blocks proestrus vs. diestrus, *p* = 5.94 × 10^−8^ and *d* = 0.48 for high blocks, two-sided Wilcoxon rank sum test *p* = 2.91 × 10^−7^ and *p* = 1.57× 10^−14^ for low and high blocks proestrus vs. males, and *p* = 0.02 and *p* = 1.54 × 10^−4^ for low and high blocks diestrus vs. males. **h.** Median block sensitivity (low - high) of detrended trial initiation times. Two-sided Wilcoxon signed-rank test for proestrus vs. diestrus *p* = 3.50 × 10^−8^ and *d* = 0.51, two-sided Wilcoxon rank sum test for proestrus vs. males *p* = 1.99 × 10^−12^ and for diestrus vs. males *p* = 1.27 × 10^−3^. **i.** Median detrended initiation times at transitions from relatively high to low (blue) or low to high value (red) blocks have greater sensitivity to the change in value in proestrus than diestrus, *N* = 118. Causal filter spanning 15 trials, bars represent *p <* 0.05 using a two-sided Wilcoxon signed-rank test. **j.** Mean regression coefficients are stronger in proestrus, one-way ANOVA *p* = 2.42 × 10^−18^ for one trial back (*d* = 0.47 for proestrus vs. diestrus), *p* = 0.85 for two trials back, *p* = 1.07 × 10^−7^ for three trials back, *p* = 9.23 × 10^−9^ for four trials back, *p* = 0.02 for five trials back, and *p* = 0.04 for six trials back). **k.** The distribution of the differences between regression coefficients of one trial back for proestrus and diestrus is significantly different from zero, one-sided Wilcoxon signed rank test *p* = 3.91 × 10^−9^. **l.** The differences in regression coefficents of one trial back for proestrus and diestrus are significantly correlated with the differences in block sensitivity for proestrus and diestrus, Pearson correlation *p* = 4.17 × 10^−6^. **m.** The differences in block sensitivity for proestrus and diestrus are significantly correlated with the change in 17*β*-estradiol levels on sampled sessions, Pearson correlation *p* = 0.01. *, *p <* 0.01, arrows are medians and error bars are all ± standard error of the mean (SEM) unless otherwise noted. **j-m** only include rats with greater initiation times in low compared to high blocks (*N* = 114 female rats in proestrus and diestrus and *N* = 179 male rats).

To manipulate their reward expectations, rats experienced uncued blocks of trials in which they were presented with low (4, 8, or 16 *µ*L) or high (16, 32, or 64*µ*L) reward volumes. These were interleaved with mixed blocks which offered all rewards (Fig. 1b). To control for potential satiety effects across the session, we regressed out initiation times against trial number. Detrended initiation times were faster in high blocks compared to low blocks (block sensitivity, Fig. 1c). Previous work has suggested that this pattern optimally balances the costs of vigor against the benefits of harvesting reward in environments with different reward rates^29^. The sensitivity of trial initiation times to the reward blocks was robust across 303 male and female rats trained on the task (Fig. 1d).

We previously found that the decision of when to initiate a trial was governed by a reinforcement learning algorithm that updated the value of the environment or state using RPEs^28^. In actor-critic models, the critic learns the value of the state (V), or the discounted sum of future expected rewards (R) in that state, and updates state values trial-by-trial by the following equation: *V_t_*_+1_ = *V_t_* + *α*(*R_t_ − V_t_*), where *R_t_ − V_t_* is the RPE (*δ*), and *α* is the learning rate (Fig. 1e). This algorithm, also called a delta rule, computes the average of previous rewards, with most recent rewards having the strongest impact. Indeed, regressing trial initiation times against previous reward offers yielded negative coefficients that declined exponentially^9^, with more recent rewards having a stronger influence on behavior (Fig. 1f). When a rat received a large reward, it initiated the next trial more quickly (i.e., rewards and initiation times were inversely proportional), consistent with positive RPEs increasing state values. The exponentially decaying influence of previous rewards on value estimates is a hallmark of reinforcement learning algorithms, including the delta rule described above. Collectively, this suggests that rats estimate their future expected rewards by integrating over past reward offers via a delta rule, and initiate trials more quickly when they anticipate higher future expected rewards.

To determine whether hormonal fluctuations modulate reinforcement learning, we tracked female rats’ reproductive or estrous cycles with vaginal cytology (Extended Data 1a). There are four stages of the estrous cycle, each lasting ∼12-48 hours. 17*β*-estradiol is low during metestrus and diestrus and peaks during proestrus, followed by estrus, which is when ovulation occurs (Extended Data 1b). Serum 17*β*-estradiol expression patterns measured with an ELISA validated stages identified by vaginal cytology (Extended Data 1c, d).

When female rats were in proestrus, the stage where 17*β*-estradiol peaks, their trial initiation times were more sensitive to the reward blocks compared to when those same rats were in diestrus, when 17*β*-estradiol is low (Fig. 1g-h, Extended Data 1e). Detrended initiation times showed stronger (weaker) modulation of initiation times following block transitions in proestrus (diestrus, Fig. 1i). The difference between trial initiation times in low and high blocks tracked 17*β*-estradiol levels across the estrous cycle, with an enhanced difference in proestrus that continued into estrus, when ovulation occurs (Extended Data 1c,e)). In diestrus, females showed block sensitivity that was closer to males, suggesting a dose-dependent effect of 17*β*-estradiol (Fig. 1g,h).

Rats also adjusted how long they were willing to wait for the 16 *µ*L rewards in the different blocks^28^. There was no effect of the estrous cycle on how long they were willing to wait for rewards (Extended Data 2). Moreover, we previously found that the wait times are governed by a different behavioral strategy that uses state inference, and that is distinct from the trial-by-trial reinforcement learning strategy that governs initiation times^28^. Because the trial initiation times reflect a reinforcement learning process that we hypothesize relies on dopaminergic RPEs, and they are hormonally modulated, we focused on that aspect of behavior in this study.

To assess whether hormonal fluctuations modulate trial-by-trial learning at a finer timescale than reward blocks, we regressed detrended trial initiation times in mixed blocks against previous rewards in proestrus and diestrus. In proestrus, detrended initiation times were more strongly influenced by the reward on the most recent trial compared to diestrus and to males (Fig. 1j, k). The increased weight of the previous reward correlated with increased behavioral sensitivity to blocks, suggesting that these metrics of learning, although at different timescales, are related (Fig. 1l). We next regressed detrended initiation times against the previous reward, but this time included stage as a categorical variable. Across rats, we found that there was a significant interaction between stage and the previous reward that increased the influence of the previous reward on behavior in proestrus (*p* = 8.43 × 10^−5^).

Notably, there was variability in the degree to which the estrous cycle modulated behavior across rats (change in block sensitivity between proestrus and diestrus, Fig. 1h). To determine the biological mechanism, we measured serum levels of 17*β*-estradiol with an ELISA in rats performing the task across the estrous cycle. Using 11 rats, we first validated that 17*β*-estradiol was indeed higher in proestrus during these training sessions and that this subset qualitatively reflected the behavioral effects we identified in the larger population (Extended Data 1d, Extended Data 3a-c). We uncovered a strong and significant correlation between changes in circulating levels of 17*β*-estradiol and hormonal modulation of block sensitivity, where rats with greater changes in 17*β*-estradiol had greater hormonal modulation of behavior (Fig. 1m). Not only does this draw a clear connection between 17*β*-estradiol and learning, but it also suggests that the degree to which 17*β*-estradiol fluctuates over the cycle dictates whether the cycle influences learning and can explain why some rats show stronger or weaker modulation.

We refrain from making claims about whether hormonal modulation of behavior, including enhanced sensitivity to previous rewards, is better or worse for performance, since the exact benefits (costs) that are being maximized (minimized) are unobservable. Instead, we used a reinforcement learning framework to make precise descriptive statements about changes in learning over hormonal states.

### 17*β*-estradiol enhances dopamine RPEs

We next sought to characterize the neural correlates of enhanced reinforcement learning in proestrus. To test whether physiological patterns of dopamine signaling encode an RPE (Fig. 2a) that is modulated by the estrous cycle, we measured dopamine in the NAcc using fiber photometry of dopamine sensors. We virally expressed the optical dopamine sensor GRAB_DA_ in the NAcc, and implanted a fiber optic over the injection site (Fig. 2b). We used a static fluorophore (mCherry) to correct for brain motion (Methods, Extended Data 5e,f). We observed a strong phasic dopamine response when the rat heard the reward offer cue. Dopamine fluorescence at the offer cue scaled with the offered reward volume: there were positive phasic responses to larger rewards and negative responses to smaller rewards, consistent with an RPE (Fig. 2c,d). We interpret this result as being consistent with a temporal difference learning algorithm^7,22^, in which the RPE would propagate back to the earliest predictor of the value of the environment on each trial (here, the cue).

**Figure 2:**
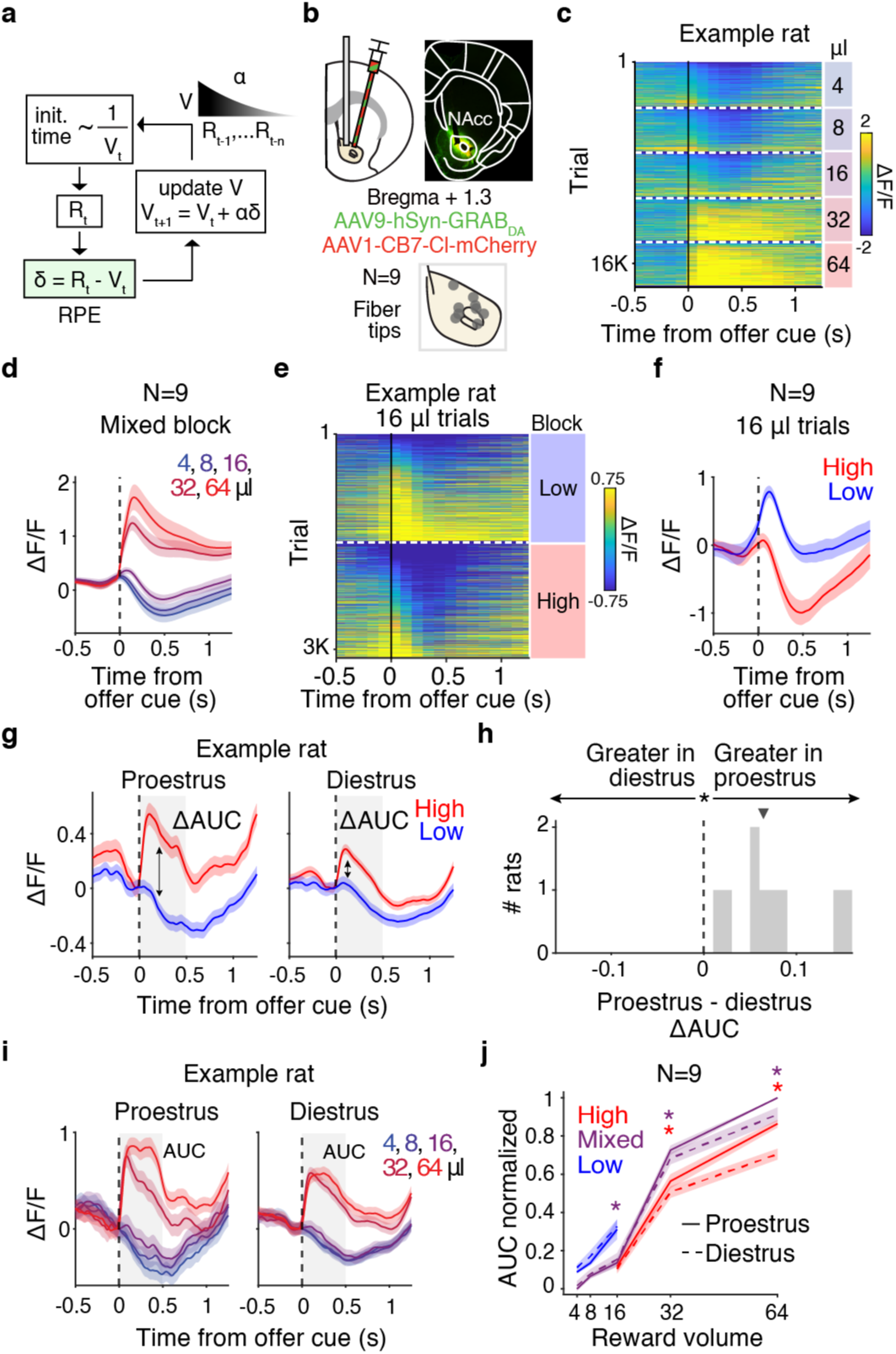
NAcc dopamine at reward offer cue encodes RPEs with greater dynamic range in proestrus. **a.** Schematic depicting delta rule with RPE highlighted. **b.** NAcc GRAB_DA_ and mCherry virus injections in female rats, with fibers. (Top left) Schematic and (top right) representative example of GRAB_DA_ expression in green and mCherry in red. (Bottom) Fiber locations in the NAcc for 9 female rats. **c.** Heatmap of GRAB_DA_ signal in response to reward offer cue across *>*16,000 trials recorded from an example rat during mixed blocks and sorted by reward volume and peak response in the first 500 ms following offer cue. Time bins are 100 ms. **d.** Average response to offer cue during mixed blocks, separated by reward volume. **e.** Heatmap of GRAB_DA_ signal in response to reward offer cue for 16 *µ*L across *>*3,000 trials recorded from an example rat sorted by reward block and peak response in the first 500 ms following offer cue. Time bins are 100ms. **f.** Average response to 16 *µ*L offer cue across the population, separated by reward block. **g.** Response to offer cue, separated by reward block and stage group for example rat. Gray box represents window used to calculate the change in area under the curve (ΔAUC) 500 ms after the offer cue in high - low blocks in **h**. **h.** Histogram of stage effect (proestrus-diestrus) on ΔAUC for offer cue response over rats shows stronger block encoding in proestrus, two-sided Wilcoxon signed-rank test *p* = 3.91 × 10^−3^. Triangle is median. **i.** Response to offer cue, separated by reward volume and stage group for an example rat. **j.** Median population response to each reward volume, separated by reward block and min-max normalized. AUC is larger in proestrus than diestrus, two-sided Wilcoxon signed-rank test during mixed blocks at 16 (*p* = 0.039, *d* = 0.19), 32 (*p* = 0.02, *d* = 0.49), and 64 (*p* = 0.0039, *d* = 0.89) *µ*L and during high blocks at 32 (*p* = 0.02, *d* = 0.53) and 64 *µ*L (*p* = 0.0039, *d* = 1.64). \**p <* 0.05, all error bars are ± SEM. In **g-j**, data is baseline-corrected using 50 ms before the offer cue.

We next examined the 16*µ*L reward offers, which appeared in all reward blocks, but with a different value relative to the average reward in each block. In reinforcement learning algorithms, the value of the state (against which rewards are compared) is the discounted sum of future expected rewards in that state. In our task, the expected rewards varied with the reward blocks. The response to the auditory cue predicting 16*µ*L was positive in low blocks, when the average reward was low, and negative in high blocks, when the average reward was high, consistent with an RPE (Fig. 2a,e,f). Therefore, we focused our subsequent analyses on the phasic dopamine response at the time of the offer cue.

We found that phasic dopamine release was more strongly modulated by reward states in proestrus, when 17*β*-estradiol was elevated, compared to diestrus (Fig. 2g,h). Similar to what was observed behaviorally, in proestrus, the phasic dopamine response to the offer cue was more sensitive to the reward blocks (Fig. 2g). To quantify this effect, for each rat, we calculated the area under the curve of the dopamine response in the 500ms following the offer cue, for low and high blocks. For all rats, there was a larger difference in these values in proestrus (Fig. 2h). Moreover, this difference was driven by enhanced dopamine transients for cues predicting the largest rewards (32 and 64 *µ*L) in proestrus (Fig. 2i,j).

Dopamine release at recording sites dorsal to the NAcc in the caudate putamen also exhibited hallmarks of RPE encoding during the offer cue (Extended Data 6). However, dopamine at these sites was unaffected by the estrous cycle, suggesting that hormonal modulation of dopamine release may be specific to the NAcc.

Previous work has found that dopamine activity in the VTA can reflect beliefs about hidden task states if rewards are probabilistic and exhibit variable timing^30^. Rewards in our task were omitted on 15-25% of trials and the delay to reward was variable. We previously found that dopamine in the NAcc exhibited similar dynamics to the VTA during the delay period when there was ambiguity about whether the rats were in a rewarded or unrewarded trial^31^. Dopamine during rewarded trials sorted by delay duration and aligned to the delay start exhibited negative ramps during the delay (Extended Data 7a) that have been interpreted as moment-by-moment negative RPEs as the animals waited without receiving a reward^12^. The magnitude of phasic dopamine aligned to the reward cue, signaling the end of delay, scaled with delay length as the reward became more surprising at longer delays (Extended Data 7b). This pattern requires additional knowledge about the task structure as the probability of being in an unrewarded trial increases over time, so the reward cue is more unexpected given beliefs about the trial^30^. Therefore, NAcc dopamine reflects state inference at the time of probabilistic rewards. Interestingly, this RPE was also larger in proestrus compared to diestrus (Extended Data 7c, d), especially at longer delays, suggesting that RPEs related to state inference were also enhanced by 17*β*-estradiol. Notably, these RPEs cannot account for the modulation of initiation times by block over estrous as the delays did not change over blocks.

To test whether enhanced RPEs at the offer cue could account for enhanced reinforcement learning in proestrus, we used a reinforcement learning model that predicts rats’ trial initiation times^28,31^. In the model, initiation times were inversely related to state value, which was estimated with a delta rule (Fig. 3a). The model captured qualitative aspects of rats’ behavior, including sensitivity of initiation times to the reward blocks and previous reward offers (Fig. 3b,c). To determine when RPEs were most strongly encoded, we fit the model to each rat’s initiation times. Then we used the best fit parameters for each rat to estimate their trial-by-trial RPEs (see Methods). We regressed RPEs from the behavioral model against the dopamine AUC in sliding windows around each event (Fig. 3d). We found the strongest encoding of RPEs (i.e., largest regression coefficients) at the time of the offer cue, when rats hear the tone indicating the reward offer, consistent with a temporal difference learning algorithm.

**Figure 3:**
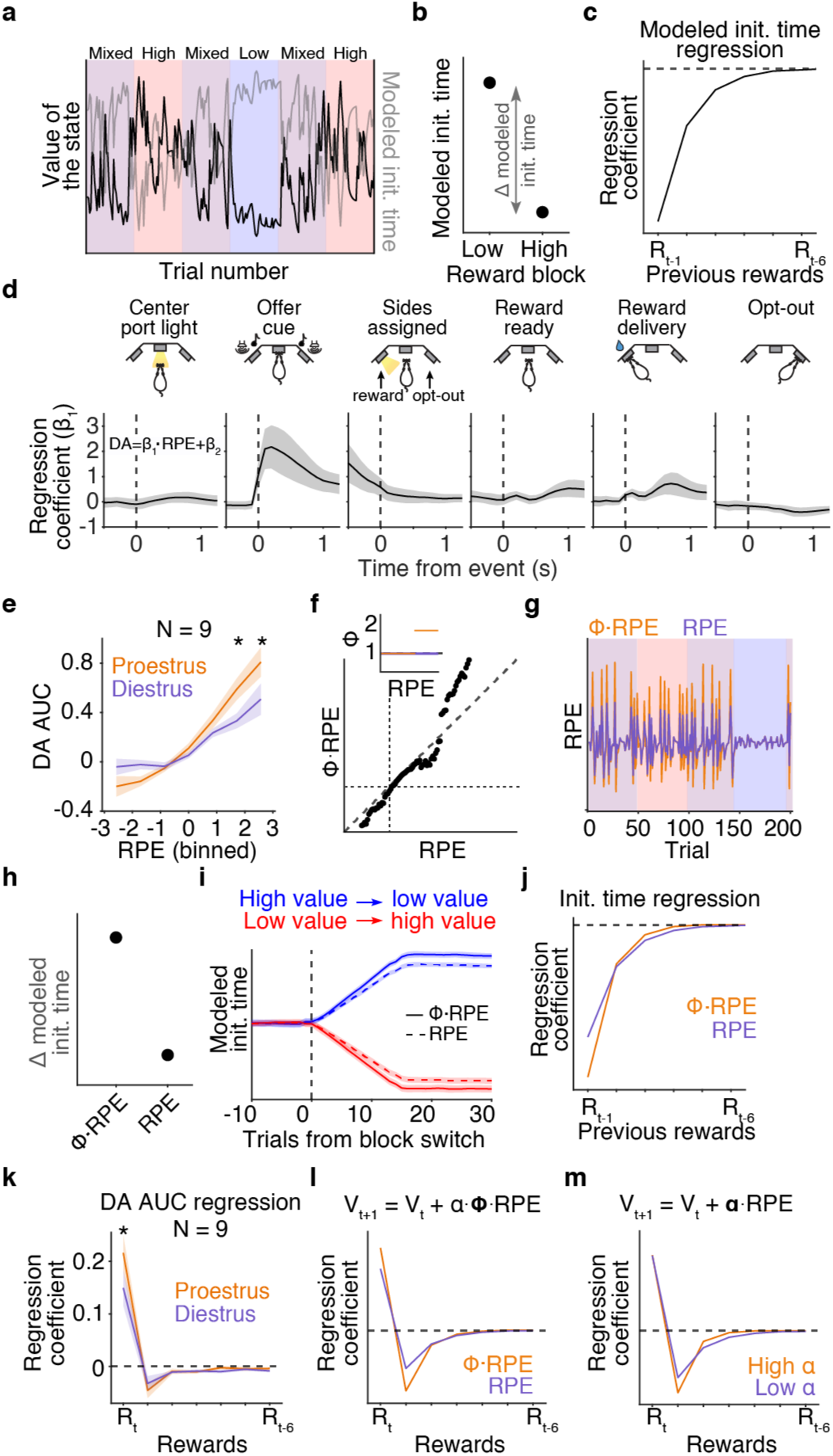
Multiplicative gain on large RPEs in a reinforcement learning model captures enhanced learning from rewards in proestrus. **a.** Simulation of the value of the state across reward blocks (blue = low, purple = mixed, red = high) using a reinforcement learning model and predicted initiation times for one session. **b.** Model-predicted initiation times by block. **c.** Regression of model-predicted initiation times against rewards on previous trials during mixed blocks. **d.** Regression coefficients of model-predicted RPE (estimated from each rat’s best fit model parameters) as a function of dopamine in each time bin (100 ms) aligned to task events, with the strongest encoding at the offer cue. **e.** Median dopamine AUC at the offer cue as a function of model-predicted RPEs is greater for proestrus than diestrus for the two largest positive RPEs (1.71 and 2.57), Wilcoxon signed-rank test *p* = 0.039, *d* = 0.64 and *p* = 0.023, *d* = 0.54. **f.** Multiplicative gain (*ϕ*, 1.75x) was applied to large RPEs (RPEs *>* 90*^th^* percentile). **g.** Increasing large RPEs enhanced trial-by-trial RPEs, mimicking proestrus. **h-j.** Increasing *ϕ* on large RPEs captures enhanced learning seen in proestrus, including greater block sensitivity (**h**), more pronounced behavioral changes at block transitions (**i**), and greater impact of previous rewards on initiation time (**j**). **k.** Rewards on the current trial have a significantly larger impact on dopamine in proestrus compared to diestrus, Wilcoxon signed-rank test *p* = 0.02, *d* = 0.21. **l-m.** Enhancing *ϕ* captured the increased regression coefficient on the current trial (**l**), while enhancing the learning rate, *α*, did not (**m**).

To quantify encoding of RPEs in proestrus and diestrus, we used the behavioral model - fit to all stages of the estrous cycle - to generate trial-by-trial RPE estimates for each rat. We then separately computed the dopamine AUC for different RPE values in proestrus versus diestrus. We found that in proestrus, dopamine encodes RPEs with a greater dynamic range. Moreover, there is a significant increase in the amplitude of the phasic dopamine response for large, positive RPEs (Fig. 3e, see also Fig. 2g,i,j).

To account for these apparent effects on large positive RPEs, we modified the behavioral model to include a multiplicative gain (*ϕ*) on positive RPEs above the *>* 90*^th^* percentile (Fig. 3f inset). This explicitly enhanced large positive RPEs; however, because these larger RPEs increased state values, this manipulation also produced a modest increase in the dynamic range of negative RPEs (Fig. 3f,g), consistent with dopamine measurements (Fig. 3e). Adding this *ϕ* parameter recapitulated the enhanced behavioral sensitivity to reward blocks in proestrus (Fig. 3h, see also Fig. 1h). Moreover, it changed the impact and time constant with which previous rewards influenced the model’s behavior, and increased behavioral modulation at block transitions, consistent with behavior in proestrus compared to diestrus (Fig. 3i,j, see also Fig. 1i,j). Alternatively, an enhanced learning rate parameter could in principle increase the impact of previous rewards on behavior. However, an increased learning rate does not predict any change in the relationship between the current reward offer and the RPE. In proestrus, dopamine showed enhanced responses to the current reward offer, which was captured by changing the *ϕ* parameter in the model, but not the learning rate, suggesting that rats in proestrus specifically exhibit larger positive RPEs (Fig. 3k-m).

Collectively, these data show that dopamine dynamics at the time of the offer cue encoded an RPE. Dopamine responses in the NAcc at this timepoint scaled with the offered reward (Fig. 2c,d), were inversely proportional to reward expectations (Fig. 2e,f), correlated with RPEs estimated from the behavioral model (Fig. 3d), and exhibited a greater dynamic range in proestrus compared to diestrus (Fig. 2h,j; Fig. 3e). Hormonal modulation of female rats’ reinforcement learning could be accounted for by a multiplicative gain on large positive RPEs, which was consistent with measurements of dopamine release in the NAcc.

### NAcc dopamine controls initiation times

According to the behavioral model, dopamine RPEs update the value of the state, which determines the vigor with which animals initiate subsequent trials. To test this hypothesis, we evaluated dopamine on trials with different subsequent initiation times. In previous work, we found that initiation times were most strongly governed by RPEs following unrewarded trials^28^, so we restricted the analysis of dopamine to those trials. Indeed, trials in which subsequent initiation times were fast showed larger phasic dopamine responses, and trials with long initiation times showed apparent dips in the dopamine signal (Fig. 4a). There was a significant negative correlation between the subsequent initiation time on trial *t* + 1, and the amplitude of the dopamine on trial *t*, in all rats (Fig. 4b,c). In other words, the phasic dopamine response at the offer cue predicted the rats’ initiation time on the next trial, consistent with dopamine at this event acting as an RPE to update the value of the state for response vigor.

**Figure 4:**
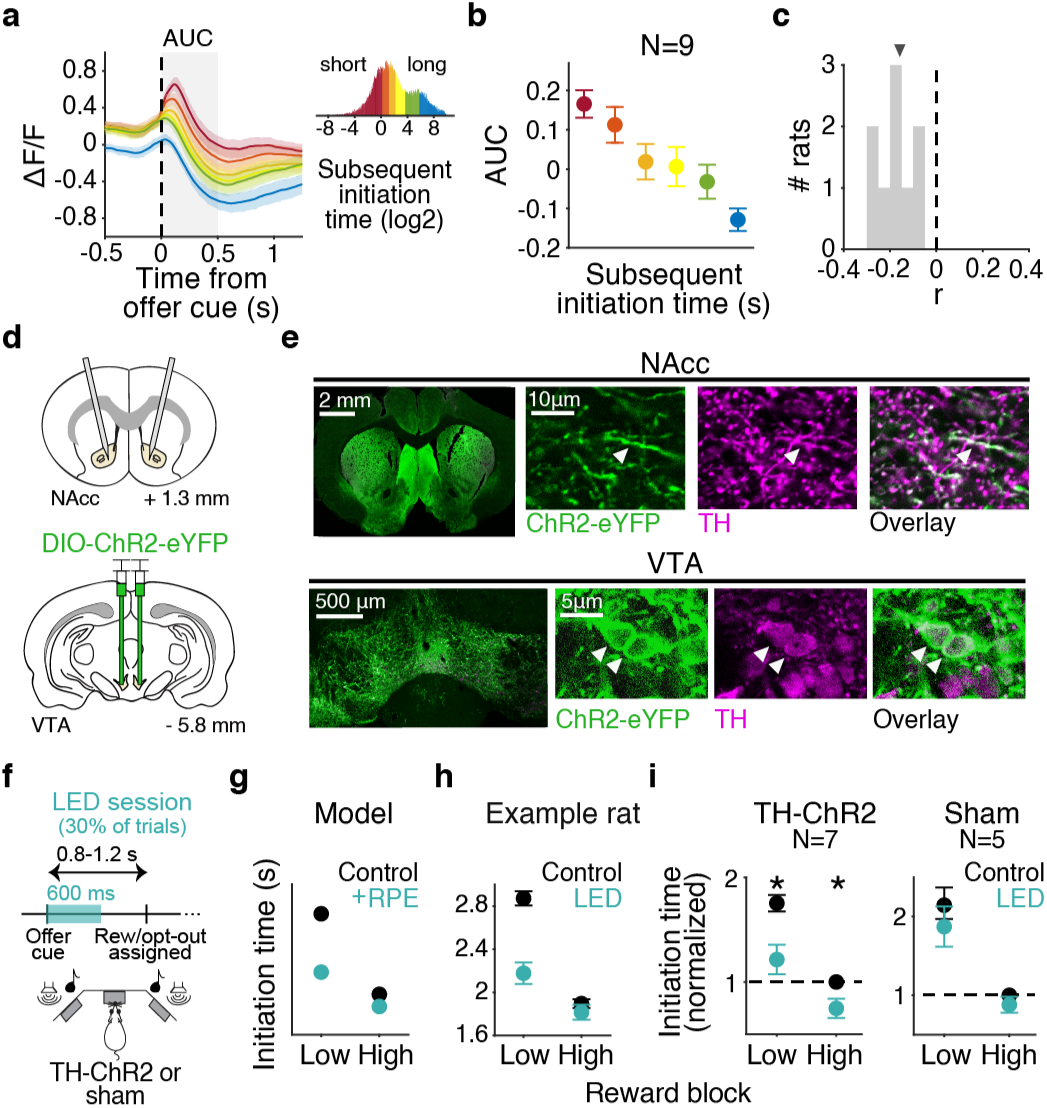
Phasic dopamine release in the NAcc controls trial initiation time, consistent with an RPE that updates state values. **a.** Dopamine signal aligned to the offer cue split by the subsequent trial initiation time (*<*0.2, 0.4, 0.9, 2.5, and 12 s and *>*12 s), *N* = 9. Inset histogram shows how log_2_(initiation times) were binned. **b.** Mean AUC of curves in 500 ms window shown in **a**. **c.** Pearson correlation of AUC and subsequent initiation time over rats, all *p <<* 0.01. Triangle is median. **d.** Schematic of the strategy used to specifically stimulate TH+ VTA terminals expressing ChR2 in NAcc. **e.** ChR2-YFP colocalization with TH in VTA cell bodies and NAcc axons (20x and 63x). Arrows point to labeled NAcc axons or VTA cell bodies. **f.** TH+ axon terminals in the NAcc were stimulated with 465 nm light during the offer cue on 30% of trials. **g.** Simulated initiation times with and without an additive RPE gain on 30% of trials, using the behavioral model. **h.** Mean trial initiation times for an example TH-Cre rat for sessions with optogenetic stimulation and without (control). **i.** Mean trial initiation times across the population normalized to high blocks in control sessions for rats expressing ChR2 in TH+ axons (left) and sham controls (right), two-sided Wilcoxon signed-rank test *p* = 0.02, *d* = 1.47 for low blocks and *p* = 0.03, *d* = 1.35 for high blocks. All error bars are mean ± SEM, \**p <* 0.05.

To test whether dopamine at this timepoint causally influenced initiation times, we used optogenetics to activate dopamine terminals in the NAcc. We expressed Cre-dependent excitatory opsins (AAV9-DIO-ChR2.0) bilaterally in the VTA and substantia nigra of tyrosine hydroxylase (TH)-Cre rats and rats injected with a TH-Cre virus, and bilaterally implanted fiber optics over the NAcc (Fig. 4d,e). We histologically verified that ChR2 expression was largely restricted to neurons that were TH-immunoreactive (Fig. 4e, Extended Data 8). On 30% of trials, we optogenetically stimulated midbrain dopamine terminals in the NAcc triggered when rats heard the tone indicating the reward offer.

We reasoned that this manipulation was similar to providing an additive gain to RPEs since it would add an (approximately) constant amount of dopamine on stimulation trials. Simulating an additive RPE gain in the behavioral model predicted that rats should exhibit faster trial initiation times because positive prediction errors would increase state values (Fig. 4g). The model predicted a larger effect in low blocks, because it implemented a floor effect to capture the fact that rats can only initiate trials so quickly in high blocks. Indeed, we found that dopamine terminal stimulation decreased trial initiation times (Fig. 4h,i). Notably, there was no effect on trial initiation times in sham animals that did not express light-sensitive opsins (Fig. 4i). There was also no effect on wait times (two-way ANOVA tests, *F* = 0.24 and *p* = 0.91 for an interaction between reward and session type for TH-Cre rats and *F* = 0.207 and *p* = 0.103 for sham rats), further showing that dopamine at reward offer cue encodes an RPE that is selectively used to guide initiation times. Collectively, these results support the hypothesis that dopamine signaling in the NAcc at the time of the reward offer acts as an RPE to adjust the value of the state for subsequent trial initiation times. We note that while we used continuous light stimulation, our results are consistent with previous optogenetic studies that used pulse-based stimulation^16,32,33^.

### Reduced dopamine reuptake in proestrus

To identify the cellular mechanisms by which reinforcement learning and RPE encoding was enhanced in the stages of the estrous cycle that are proximal to the peak in 17*β*-estradiol (proestrus, and to a lesser extent, estrus, Extended Data 1e; Extended Data 5a-d), we performed quantitative mass spectrometry of the NAcc in diestrus, proestrus, and estrus (*N* = 5-6 rats in each group). 238 proteins were significantly decreased in expression in proestrus or estrus compared to diestrus, and 258 increased. Gene ontology analysis identified several proteins involved in dopamine signaling that were differentially expressed in these stages (Methods, Extended Data 9). However, the only differentially expressed proteins that could account for the observed increase in phasic dopamine in proestrus and estrus were proteins involved in dopamine reuptake, which were expressed at lower levels in these stages (Fig. 5a, Extended Data 9). The dopamine transporter (DAT or Slc6a3) controls dopamine reuptake in the NAcc, and was expressed at significantly lower levels in proestrus compared to diestrus. The serotonin transporter (SERT or Slc6a4) is also capable of robust dopamine reuptake^34^, and was expressed at lower levels in proestrus and estrus compared to diestrus. Smpd3 (sphingomyeline phosphodiesterase 3) triggers fusion of DAT to the plasma membrane, and has been shown to make dopamine reuptake more efficacious^35,36^. Smpd3 was decreased in expression in estrus, suggesting a lower density of DAT in the plasma membrane during estrus, consistent with effects of 17*β*-estradiol applications on DAT localization in vitro^36^. Reduced plasmalemmal DAT and SERT should in principle lead to higher levels of extracellular dopamine.

**Figure 5:**
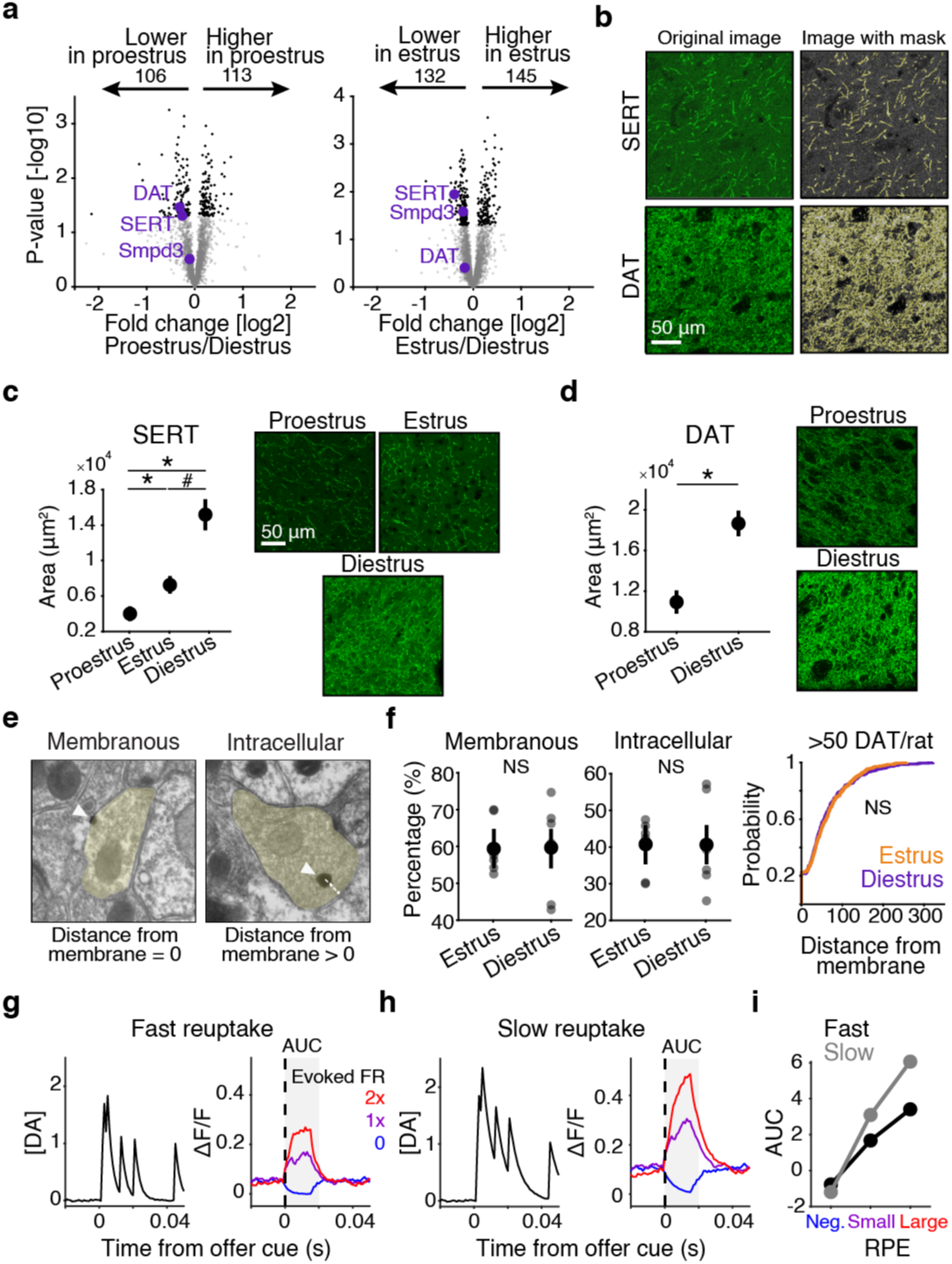
Dopamine reuptake proteins are decreased in expression in proestrus and estrus. **a.** Volcano plot depicting significantly differentially expressed proteins (with FDR correction of 5%) in black. Proteins related to dopamine reuptake that were decreased in expression in proestrus and estrus are highlighted in purple. **b.** Example images from immunohistochemical quantification of SERT and DAT expression at 63x with overlaid mask identifying fluorescent pixels above a threshold. **c.** Area of mask segmentation for SERT was highest in diestrus, Kruskal-Wallis test for group effect *p* = 4.89 × 10^−3^ and Wilcoxon rank sum post-hoc tests *p* = 5.17 × 10^−3^, *d* = 2.2 for proestrus vs. diestrus, *p* = 0.052, *d* = 1.13 for estrus vs. diestrus, and *p* = 0.024, *d* = 0.64 for proestrus vs. estrus. Example images from each stage. **d.** DAT was also reduced in proestrus compared to diestrus, two-sided Wilcoxon rank sum test *p* = 3.66 × 10^−5^, *d* = 0.92. **e.** Example of DAT labeled with silver-intensified gold and imaged with electron microscopy. Arrows point to putative DAT proteins. Minimum distance between plasma membrane and center of DAT particle are shown with dotted lines. Axons are highlighted in yellow. **f.** Proportion of membranous and intracellular DAT particles and their distance from the membrane, expressed as a cumulative probability, quantified from electron microscopy images does not change by stage. **g-h.** Simulation of dopamine concentrations and fluorescence following different evoked firing rates (FR), with fast **(g)** or slow **(h)** reuptake dynamics. **i.** Quantification of AUC in **g-h**. All error bars are mean ± SEM, \**p <* 0.05, #*p <* 0.1.

We verified that rats exhibited reduced expression levels of DAT and SERT in proestrus and estrus, using a different assay, immunohistochemistry, and a separate cohort of rats in diestrus, proestrus, and estrus (*N* = 3 per group). All sections were co-incubated in the same solutions simultaneously, to ensure that differences in fluorescence did not reflect subtle differences in tissue processing. We quantified fluorescently labeled pixel area from confocal photomicrographs, where fluorescence indicated DAT or SERT immunoreactivity (Fig. 5b). Consistent with the proteomics results of reduced DAT and SERT expression in these stages, we found that significantly fewer pixels were DAT- and SERT-immunoreactive in proestrus and estrus compared to diestrus (Fig. 5c-d). It is notable that these results held across different assays (proteomics and immunohistochemistry) and cohorts of rats.

Smpd3 is implicated in triggering fusion of DAT to the plasma membrane, and was also decreased in expression in estrus. To test whether there was also a change in subcellular localization of DAT in estrus, we characterized DAT localization at the ultrastructural level. Individual DAT particles were labeled with silver-intensified gold and visualized with electron microscopy. We quantified the fraction of DAT particles that were continuous with the membrane (membranous) versus discontinuous (intracellular), and also measured the distance from the center of the intensified gold particle to the plasma membrane. There was no significant difference in the fraction of DAT particles that were membranous versus intracellular, nor was there a difference in DAT particles’ proximity to the membrane between estrus and diestrus (Fig. 5e,f). Therefore, reduced levels of DAT and SERT expression, but not differences in subcellular localization of DAT likely accounted for enhanced dopamine RPEs and learning in proestrus (and to a lesser extent, estrus) compared to diestrus.

How could changes in reuptake, which ostensibly would result in elevated levels of extracellular dopamine, account for the enhanced phasic responses we observed? We created a computational model in which action potentials of dopamine neurons caused instantaneous increases in dopamine concentration, which decayed with a rate that reflected reuptake dynamics, unless another action potential caused another instantaneous increase. The fluorescent signal at each time step was treated as a noisy observation of the underlying dopamine concentration^37^. The model assumed that the Poisson spike rate of the dopamine neurons increased or decreased (to zero) for positive and negative RPEs, respectively. When the dopamine concentration decayed more slowly, to simulate reduced reuptake, this increased the likelihood that dopamine from consecutive action potentials would temporally summate before decaying back to baseline. Averaged over many trials, this temporal summation mechanism was sufficient to produce larger positive phasic responses (Fig. 5g-i), consistent with what we observed by measuring dopamine in proestrus at the offer cue (Fig. 2j; Fig. 3e) and after the reward delay (Extended Data 7d). Moreover, increasing temporal summation modestly elevated baseline levels of dopamine, producing a greater dynamic range for negative RPEs, which is normally thought to be limited by the low background firing rates of dopamine neurons^9^. These results are consistent with previous studies showing that inhibition of DAT increases both tonic and phasic dopamine^38,39^. In our simulation, reduced reuptake produced the strongest enhancement of the largest phasic DA responses, similar to our empirical observation that large positive RPEs were most strongly enhanced in proestrus. Therefore, decreased expression of DAT could produce a greater dynamic range of phasic dopaminergic RPEs by promoting temporal summation of dopamine transients from consecutive action potentials.

### Midbrain ER*α* promotes learning

Many hormones fluctuate over the estrous cycle. We next sought to test whether estrogenic signaling in the midbrain, specifically, influenced reinforcement learning. Expression of DAT in the NAcc is restricted to dopaminergic axons from the midbrain. Therefore, we hypothesized that estrogen receptors, which can regulate protein expression via transcription and other mechanisms when bound to 17*β*-estradiol^1,17,18^, might modulate learning by influencing expression of DAT in dopaminergic neurons. We virally expressed short-hairpin RNA (shRNA), labeled with eYFP, to knockdown expression of the estrogen receptor ER*α* bilaterally in the VTA (shEsr1-eYFP, Fig. 6a,b). We used either a virus that was inducible by doxycycline in food and water (U6/tetO-shEsr1) or one that was constitutively active (lenti-shEsr1, Methods). For each virus, we used fluorescent in situ hybridization of fixed tissue sections to measure levels of labeled *Esr1* (ER*α* transcript). Quantification of labeled probes showed a significant reduction in *Esr1* in rats virally expressing shRNAs (Fig. 6c,d).

**Figure 6:**
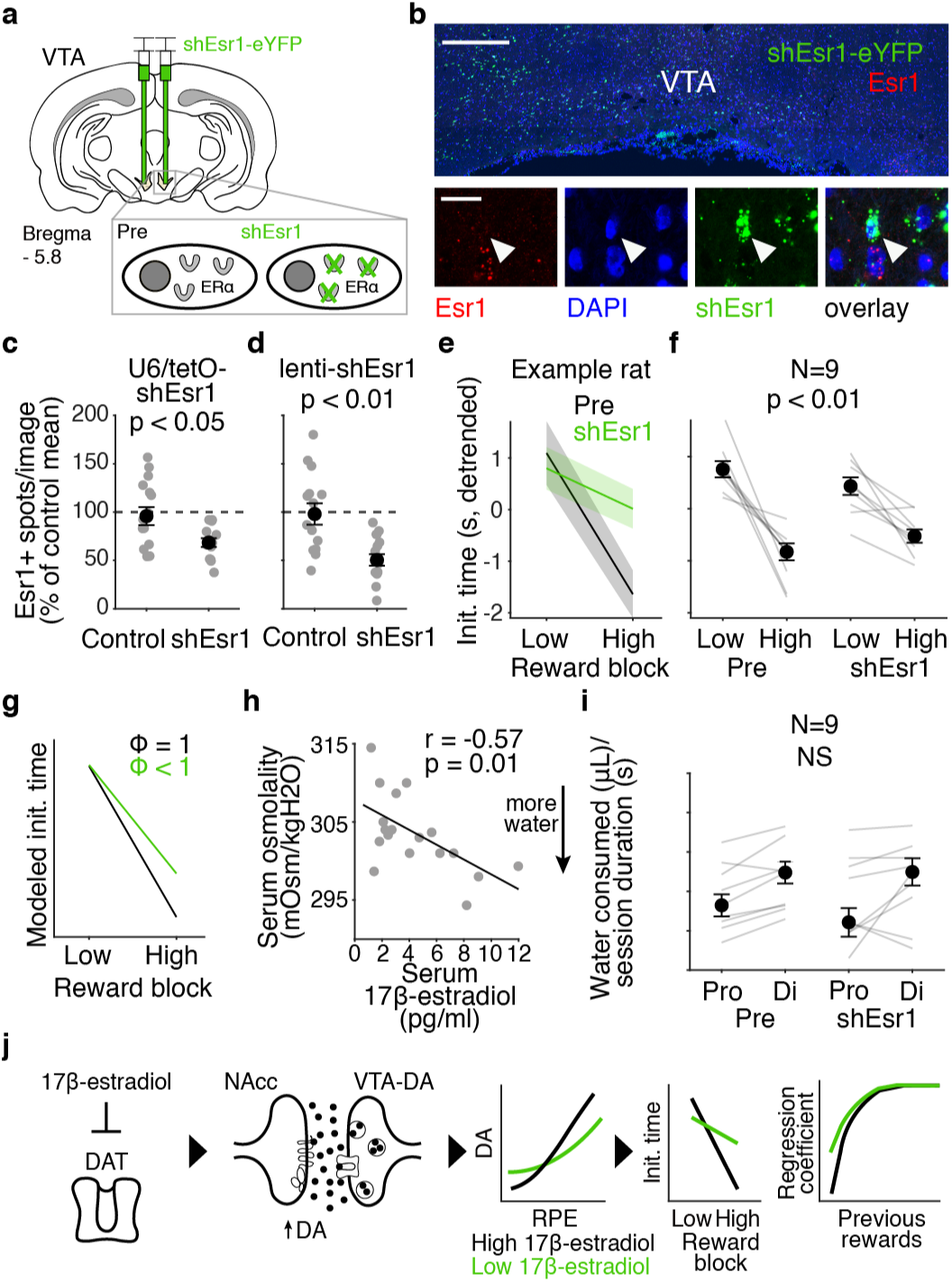
Midbrain estrogen receptor (ER*α*) knockdown suppresses reinforcement learning. **a.** shEsr1-eYFP was injected bilaterally in the VTA. **b.** Expression of eYFP-labeled shEsr1 virus in the VTA (20x, top) after RNAscope. Scale bar is 500 *µ*m. Expression of *Esr1* in eYFP-labeled cells compared to cells not expressing eYFP (63x, bottom). Scale bar is 20 *µ*m. Arrow points to lack of *Esr1* expression in eYFP-labeled cell. **c-d.** Number of *Esr1*-labeled spots per photomicrograph of a z-stack taken from a rat injected (**c**) with U6/tetO-shEsr1 and (**d**) lenti-shEsr1 compared to not injected control tissue, expressed as a percentage of the control means. *Esr1* is suppressed with shEsr1 compared to controls, two-sided Wilcoxon rank sum tests for U6/tetO-shEsr1 *p* = 0.014, *d* = 1.0 and lenti-shEsr1 *p* = 2.41 × 10^−3^, *d* = 1.15. **e.** Mean detrended initiation time of one example rat before and after shEsr1 injection for low and high blocks. **f.** Detrended initiation time across the population before and after shEsr1 injection for low and high blocks. Block sensitivity (low - high) was significantly reduced after shEsr1, two-sided Wilcoxon signed rank test *p* = 3.91 × 10^−3^, *d* = 1.12. **g.** Reducing *ϕ* to below 1 replicates suppression of behavioral sensitivity with shEsr1. **h.** Serum osmolality is significantly correlated with serum 17*β*-estradiol. Line of best fit overlaid (*N* = 18). **i.** Water consumed per session, controlling for the total amount of time the rat was allotted in the task, was still reduced in proestrus (Pro) compared to diestrus (Di) and this reduction was not affected by shEsr1, two-sided Wilcoxon signed rank test comparing difference in water consumption in proestrus and diestrus before injection to after shEsr1 *p* = 0.82 and *d* = 0.031. **j.** Schematic summarizing findings. 17*β*-estradiol reduces expression of dopamine transporters. This increases the amount of dopamine in the extracellular space, allowing for a greater dynamic range of RPEs and enhanced learning from previous rewards in proestrus. All error bars are median ± SEM, unless otherwise noted.

Genetic suppression of estrogen receptors in the midbrain significantly reduced rats’ behavioral sensitivity to the reward blocks (Fig. 6e,f). This is consistent with the finding that block sensitivity was correlated with 17*β*-estradiol over the estrous cycle, suggesting a dose-dependent effect of 17*β*-estradiol on block sensitivity. Reducing the RPE gain parameter in the delta model recapitulated the effect of shEsr1 (6g). We conclude that midbrain estrogen receptors promote reinforcement learning by increasing the impact of previous rewards on behavior.

In proestrus, rats also appeared to be less thirsty: their initiation times were significantly slower, they performed fewer behavioral trials and drank less water, compared to diestrus (Extended Data 10a-c). Measurements of serum osmolality indicated reduced thirst in proestrus, and were strongly correlated with serum levels of 17*β*-estradiol (Extended Data 10d-e; Fig. 6h). Notably, detrending the initiation times minimized satiety effects, and allowed us to resolve estrogenic influences on trial-by-trial reinforcement learning. However, because the ER*α* knockdown was specific to the VTA, this experiment provided an opportunity to directly test whether estrogenic modulation of reinforcement learning occurs independent of systemic effects on fluid balance. We compared rats’ water consumption before and after knockdown of ER*α*. Despite reducing behavioral sensitivity to the reward blocks, ER*α* knockdown had no effect on estrous-dependent water consumption (6i). These results indicate that 17*β*-estradiol impacts reinforcement learning and fluid balance through dissociable mechanisms: the former relies on ER*α* in the VTA, but the latter does not.

## Discussion

Using a task that tested reinforcement learning, we found that rats learn more from previous rewards when they express higher levels of 17*β*-estradiol, building on previous studies that found sex-dependent effects of reward on motivation^40^. We reveal a computational and circuit-level mechanism that underlies midbrain 17*β*-estradiol-dependent enhanced learning: expression of the dopamine transporter in the NAcc is reduced in proestrus, resulting in a larger dynamic range of dopamine levels that enhances RPEs, which causally influence initiation times on subsequent trials (Fig. 6j). Our findings suggest that the enhancement is most pronounced for large positive RPEs, possibly due to increased occupancy of pools of DAT proteins at high dopamine concentrations. Expression of DAT and SERT in the striatum is highly dynamic^41,42^, with protein turnover occurring on the order of hours to days^42–44^, suggesting that hormonal fluctuations over estrous could plausibly affect plasmalemmal transporter expression.

One key future question is whether 17*β*-estradiol also modulates the firing rates of dopaminergic cells in the VTA, in addition to dopamine dynamics in the NAcc. While previous studies have examined estrous-dependent VTA dynamics in the context of drug exposure^45,46^, it is unclear whether 17*β*-estradiol affects encoding of RPEs during reinforcement learning at the level of spikes in the VTA. 17*β*-estradiol could in principle act directly on VTA neurons^47,48^, or have indirect effects by modulating the excitability of afferent inputs to the VTA^49^. Indeed, medium spiny neurons in the NAcc are known to project back to the VTA^50^, so it is possible that the estrogenic modulation of dopamine dynamics in the NAcc described here could indirectly influence VTA activity, via this projection. Distinct subpopulations of dopamine neurons in the VTA broadcast RPEs to different downstream circuits^51^, so a question for future exploration is whether estrous-dependent modulation of RPEs is circuit-specific, and largely restricted to state value learning in the NAcc, or includes the dorsal striatum. We found that dopamine release at sites that were dorsal to the NAcc were not modulated by estrous, suggesting regional specificity of estrogenic modulation. Local modulation of reuptake may provide more spatially precise control of dopamine signaling compared to regulation at the level of somatically-generated action potentials, given the enormous axonal arborizations of dopamine neurons^52^.

Here, we find a novel role for dopamine transporters in shaping RPEs for reinforcement learning. Drugs of abuse such as cocaine and methamphetamine act on DAT^53^, and both DAT and SERT are the targets of pharmacological treatments for many neuropsychiatric disorders, including schizophrenia, depression, anxiety, and attention deficit hyperactivity disorder^54^. Computational psychiatry models have examined these disorders through the lens of reinforcement learning^55–57^. Our results provide a potential biological mechanism that bridges dopamine reuptake and reinforcement learning in health and disease, and might also underlie hormonal modulation of symptom severity in females^2–4^. The mechanisms by which ER*α* activation leads to reduced expression of dopamine reuptake proteins are important future directions. These could involve ER*α*-induced signal transduction pathways that control transcription, translation, or protein degradation, or a mechanism involving other hormone receptors such as the progesterone receptor, a canonical target of ER*α*^58^.

There is a major gap in our understanding of how endogenous hormones interact with neuromodulatory systems on fast timescales during rich, quantifiable behaviors. We developed a reinforcement learning model that accounted for rats’ behavior, and that included a computation that dopamine is thought to instantiate in the brain. Our computational and decision-theoretic approach provided a principled framework for interpreting behavioral, physiological, optogenetic, and molecular profiling data, and represents a novel approach for studying endocrine systems in the brain. This approach allowed us to characterize estrogenic modulation of dopamine at broad levels of description, and opens up new avenues of research of hormone-neuromodulatory interactions during complex cognitive behaviors.

## Methods

### Subjects

A total of 365 Long-evans rats (190 male, 175 female) between the ages of 6 and 24 months were used for this study (*Rattus norvegicus*), including TH-Cre (*N* = 23), ADORA2A-Cre (*N* = 10), and DRD1-Cre (*N* = 3). Animal use procedures were approved by the New York University Animal Welfare Committee (UAWC #2021-1120) and carried out in accordance with National Institutes of Health standards.

Rats were pair-housed when possible, and water restricted to motivate them to perform behavioral trials. From Monday to Friday, they obtained water during behavioral training sessions, which were typically 90 minutes per day, and a subsequent ad libitum period of ∼ 20 minutes. Following training on Friday until mid-day Sunday, they received ad libitum water. Rats were weighed daily.

### Behavioral training

We have previously published a detailed description of behavioral training for this task^28^. Reward blocks were 40 completed trials, in which the rat remained nose poking in the center port for a variable period randomly drawn from a uniform distribution from [0.8, 1.2] seconds. On each trial, an auditory tone [1, 2, 4, 8, 16kHz] signaled one of the five different rewards. There was a one-to-one mapping between tones and reward volumes, from smallest to largest. On each trial, the reward delay period was randomly drawn from an exponential distribution with a mean of 2.5 s; the catch probability was 15-35% until the rat matched the catch rate with opt-out rate, and then lowered to 15-25%.

Early cohorts of female rats experienced the same reward set as the males [5, 10, 20, 40, 80*µ*L]. However, female rats consumed less water due to their smaller body size and performed substantially fewer trials than the males. Reward offers for female rats were slightly reduced while maintaining the logarithmic spacing to [4, 8, 16, 32, 64 *µ*L] to obtain a sufficient number of behavioral trials. For analysis, reward volumes were treated as equivalent to the corresponding volume for the male rats.

#### Criteria for including behavioral data

To determine when rats were sufficiently trained to understand the mapping between the auditory cues and water rewards, we evaluated their wait times on catch trials as a function of offered rewards. Only sessions with a running average slope (using a window of 5 sessions) above 0.01 (the average slope of shuffled data) were included to accommodate cycle-dependent changes in task performance. If fewer than three sessions per stage were included in the analysis, then this criteria was not imposed. For the comparison of water consumed over the estrous cycle (Extended Data 10), we wanted to use the same sessions for all plots, so we imposed these criteria for behavioral performance. For the shEsr1 experiment, screening for behavioral performance did not provide sufficient sessions for the analysis of water consumption (Fig. 6i). Therefore, for this analysis, we included all sessions when analyzing water consumption. We emphasize that the criteria for including sessions did not evaluate rats’ sensitivity to the reward blocks, or their trial initiation times.

#### Estrous cycle tracking

We tracked the estrous cycle for all females included in this study using vaginal cytology, with vaginal swabs collected immediately after each session using a cotton-tipped applicator first dipped in de-ionized water. Samples were smeared onto a clean glass slide and visually classified under bright-field illumination on a Leica ICC50 E microscope at 10X. The estrous cycle stage on each day was manually determined on the basis of the proportion of leukocytes, cornified epithelial cells, and nucleated epithelial cells according to established protocols using unstained smears^59^. Given that experiments were performed over the course of months, days to weeks of irregular cycling were captured and included in the data presented. Once the rat entered reproductive senescence, determined by a continuous estrus phase lasting longer than two weeks, all sessions during that time were excluded from the analyses.

To validate the accuracy of our visual classification, we measured 17*β*-estradiol in serum from trunk blood, collected in serum-separating tubes and spun at 3,000 rpm at 4°C for 10 min, with the mouse/rat estradiol ELISA kit from Calbiotech (ES180S-100) after determining stage with the method described above in 18 rats (4 in proestrus *>* 5 hours before lights out; 4 in proestrus within 3 hours before lights out; 5 in estrus; 5 in diestrus), run in triplicates. Behavioral data was collected from proestrus rats during the window when their 17*β*-estradiol was elevated (i.e. within 3 hours before lights out). For this ELISA Kit, the functional sensitivity (lowest concentration with a coefficient of variation (%CV) *<*20%) was 3 pg/ml, the precision was 3.1% (intra-assay) and 9.9% (inter-assay), and the cross-reactivity with other endogenous steroids was 0.0002% with progesterone, 0.0001% with androstenedione, 0.0002% with testosterone, and 0.0001% with cortisol^60^. All other endogenous steroids tested were *<*0.0001% or undetectable.

We measured serum 17*β*-estradiol in behaving rats using the Rat Estradiol ELISA kit from Cusabio (CSB-E05110r). For this kit, the minimum detectable amount of rat estradiol was typically less than 40 pg/ml. No significant cross-reactivity or interference between rat estradiol and analogues was observed. The intra-assay precision was CV%*<* 15% and the inter-assay precision was CV%*<*15%. For each rat, ∼ 200*µ*L blood was drawn from the lateral saphenous vein twice a week and proestrus and diestrus were sampled three times each.

#### Behavioral model

When fitting the model to rats’ initiation times (e.g., to estimate reward prediction errors), we modeled the trial initiation times as previously described (see [31] for details). Briefly, we modeled initiation times as being negatively correlated to state value according to the following equation^29,31^:

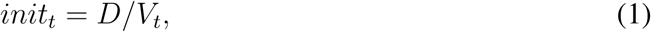

where *V_t_* represents the recency-weighted average of all previous rewards, and *D* is a scaling factor. The estimate of average reward depends on the learning rate parameter *α* with the recursive equation

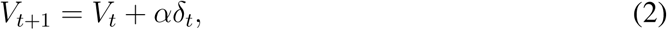

where *V_t_* is the value of the state on trial *t*, *r_t_* is the reward on trial *t*, *δ_t_* = *r_t_ − V_t_* is the RPE, and *α* is a learning rate between 0 and 1. Because we previously found that rats exhibited a dynamic learning rate^31^, we restricted model fitting to the later portion of mixed blocks, when learning rates were stable, and only related dopamine to reward prediction errors on late mixed block trials.

For this model, the activation function that converts values into a behavioral policy was a reciprocal function, which was chosen based on previous literature^28,29^, and our finding that we could reliably recover generative parameters from this model^31^. However, when simulating this model, because the reciprocal function is nonlinear, the results were sensitive to choices of other parameters (e.g., the scaling factor). Therefore, to improve the interpretability of the model simulations and make them more robust to choices about parameter values, model simulations used a linear activation function, in which the initiation times were negatively correlated with state values, parameterized with an offset and slope parameter. All model simulations used a learning rate parameter of 0.6.

#### Dopamine reuptake model

We modeled the activity of dopamine neurons as Poisson spikes with a background firing rate that increased or decreased (to zero) to reflect positive or negative RPEs, respectively. The dopamine concentration exhibited instantaneous increases at action potentials, which decayed over time to reflect reuptake, according to the following equation:

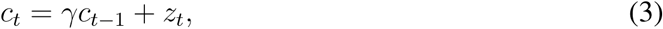

where *c_t_* is the dopamine concentration, *γ* determines the rate of decay or reuptake, and *z_t_* = 1 indicates the presence of a spike at time t, or 0 otherwise. The measured dopamine fluorescence at each timepoint, *y_t_*, was treated as an observation of the true concentration with Gaussian noise:

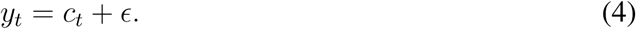

We simulated 500 trials for each condition (positive and negative RPEs), and averaged the measured dopamine signal over those trials. We performed this simulation for *γ* = 0.65, corresponding to faster reuptake, and *γ* = 0.85, corresponding to slower reuptake.

#### Stereotaxic surgeries

All surgeries were performed in female rats after 4 months of age with a Neurostar Robot Stereotaxic, using 3% isofluorane in oxygen at a flow rate of 2.5 L/minute for induction and 2% isofluorane in oxygen at a flow rate of 1.75 L/minute as maintenance for the duration of the procedure. VTA injections were conducted at a 10° angle from the midline and targeted AP -5.8; ML 0.4-0.6; DV -7.8-8.3. NAcc injections and implants targeted AP 1.3; ML 1.65; DV -6.9 with an 8-10° angle from the midline for bilateral implants.

#### Photometry

We used fiber photometry and the GRAB_DA_ dopamine sensor (AddGene #140554) to measure dopamine activity during the task. AAV9-hsyn-GRAB DA2h was injected into the NAcc. To control for motion artifacts, mCherry (AAV1-CB7-CI-mCherry-WPRE-RBG, AddGene #105-544) was also expressed because it is constitutively active. 60 nl of equal parts GRAB_DA_ and mCherry were delivered in each injection. Chronically implantable optic fibers (Thor labs) with 400 *µ*m core, 0.5 NA fiber optics were implanted unilaterally over the injection site (DV -6.7). Doric Lenses hardware and software (Doric Neuroscience Studio) were used to record fluorescence. Two-channel motion artifact correction (TMAC) was used to correct for movement artifacts, with mCherry as the activity-independent channel^61^.

#### Optogenetics

We photoactivated VTA axons in NAcc by bilaterally expressing AAV9-EF1a-double floxed-hChR2 (AddGene #20298), 500 nl per injection, in VTA of TH-Cre rats (*N* = 5) or simultaneously expressing AAV9-rTH-PI-Cre (AddGene #107788, *N* = 2, 250 nl ChR2 and 250 nl TH-Cre) to restrict expression to dopaminergic projections. Two tapered fibers (0.66 NA, 1 mm emitting length, 11 mm implant length, 200 *µ*m core diameter, Optogenix) were implanted in stimulation and sham rats as controls (*N* = 5). Continuous blue light (465 nm PlexBright LEDs, Plexon) was delivered randomly on 30% of trials for 500-700 ms during the center poke, reaching 7.5 mA.

#### Relative protein abundance profiling

16 rats (5 in diestrus, 5 in proestrus, and 6 in estrus), anesthetized with isofluorane, were decapitated and a brain matrix (Braintree Scientific) was used to make 1.0 mm coronal brain slices containing the central aspects of the NAcc (∼ 0–1.6 anterior to Bregma), kept on wet and dry ice. Bilateral 15 gauge punches of the NAcc were obtained and immediately placed on dry ice. Samples were reduced with dithiothreitol at 57° C for 1 hour (5 *µ*l of 0.2 M), alkylated with iodoacetamide at RT in the dark for 45 minutes (5 *µ*l of 0.5 M). Urea concentration was lowered to *<*2 M by addition of 100 mM Ammonium Bicarbonate and sequencing-grade modified trypsin 1.5 *µ*g (Promega) was added to each sample. Digestion proceeded overnight on a shaker at RT. To extract proteins, samples were de-salted with Sep-Pak C18 Cartridges. C18 Cartridges were conditioned by passing 1 ml of 90% acetonitrile in 0.5% acetic acid solution three times, then they were equilibrated by passing 3 ml of 0.1% TFA solution through them two times. Acidified samples were slowly loaded onto corresponding C18 cartridge packing. Samples were de-salted by passing 1 ml of 0.1% TFA solution through the cartridge three times. Then the cartridge was washed with 1 ml of 0.5% acetic acid. Peptides were eluted into clean tubes by slowly passing 500 *µ*l of 40% acetonitrile in 0.5% acetic acid, followed by the addition of 80% acetonitrile in 0.5% acetic acid.

Samples were reconstituted in 30 *µ*l of 100 mM HEPES solution, pH ∼8.5 and labeled with TMT dissolved in 90 *µ*l of 100% anhydrous EtOH for 5 min. Each sample was combined with its corresponding TMT label and incubated for 1 hour. The reaction was quenched by adding 50 *µ*l of 100 mM ammonium bicarbonate solution and incubating for 15 min. Samples were dried in a SpeedVac concentrator. The sample was then fractionated into 45 fractions using basic HPLC fractionation and the fractions were concatenated into 15 final fractions for LC-MS analysis where 1*/*10*^th^* of the fractions were analyzed individually. LC separation online with MS using the autosampler of an EASY-nLC 1000 (Thermo Scientific). Peptides were gradient eluted from the column directly to Eclipse mass spectrometer using a 1 hour gradient (Thermo Scientific) using 2% acetonitrile and 0.5% acetic acid for solvent A and 80% acetonitrile and 0.5% acetic acid for solvent B.

High resolution full MS spectra were acquired with a resolution of 60,000, an AGC target of 4e5, with a maximum ion time of 50 ms, and scan range of 400 to 1500 m/z. Following each full MS top ten data dependent HCD MS/MS spectra were acquired. All MS/MS spectra were collected using the following instrument parameters: resolution of 60,000, AGC target of 1e5, maximum ion time of 60 ms, one microscan, 0.7 m/z isolation window, and NCE of 30. MS/MS spectra were searched against the rat database using MaxQuant.

Only proteins that had at least two peptides detected and that were identified in at least three samples of one estrous stage were included in the analysis. 3082 proteins were identified and 3043 were quantified. Reporter ion intensity values were normalized and all protein intensities were Log2 transformed.

#### Electron microscopy

Rats (*N* = 12) were anesthetized with isofluorane and perfused through the ascending aorta with 50 mL of PBS containing 0.2 mL of heparin and then 4% paraformaldehyde dissolved in 0.1M phosphate buffer, pH 7.4. The brains were removed from the calvariae and post-fixed for one week in 4% paraformaldehyde. Sections through the NAcc (50 *µ*m) were cut with a vibratome, and stored in 0.01 M PBS and 0.05% azide. To enhance the antibody’s penetration of tissue, free-floating vibratome sections underwent a freeze/thaw protocol, for which tissue was cryoprotected in increasing concentrations of dimethyl sulfoxide (DMSO; 5%, 10%, and 20%) for 10 minutes each before rapid freezing in a beaker of 4-methylbutane chilled using dry ice and 100% ethanol (EtOH) as the heat conduit. After rapid freezing, the tissue was thawed by immersing in a bath of 20% DMSO at RT, before another bout of rapid freezing for a total of 8 freeze/thaw sequences. Tissue was then incubated in decreasing concentrations of DMSO (10% and 5%) for 5 minutes each before returning to PBS^62^. Tissue was stained for DAT using the protocol described above, except with an anti-DAT concentration of 1:250 that the tissue was treated with for 48 hours. Secondary antibody incubation was overnight in PBS-BSA-Azide containing ultra-small colloidal gold-conjugated goat anti-rabbit secondary antibody (1:100, EMSciences #25181, lot #GAR-01217/1). Sections were rinsed in PBS and post-fixed in 2% glutaraldehyde in PBS for 10 min, and then the steps for silver intensification (KPL Silver Enhancer Kit, #5520-0021) were carried out to enlarge the ultra-small colloidal gold particles.

The EM tissue processing was similar to the procedure performed before^63^, which consisted of osmium-free tissue processing in 1% tannic acid, 1% uranyl acetate and 0.2% iridium tetrabromide, each dissolved in maleate buffer (pH 6.0, 0.1 M), post-fixation in 1% uranyl acetate in 70% EtOH overnight, dehydration in 70%, 90%, and 100% EtOH, followed by 100% acetone, infiltration in EPON 812 (EM Sciences), flat-embed between Aclar plastic sheets (EMSciences) and capsule-embedded in BEEM capsules (EMSciences). Ultrathin sections were prepared using the Ultracut E ultratomicrotome and collected on nickel grids (EMSciences). Sections were counter-stained with lead citrate to improve contrast. Images were captured at a magnification of 50,000× using the Hamamatsu CCD camera (1.2 megapixel) and software from AMT (Boston, MA, USA).

For each animal, approximately 50 axons were sampled and quantified. Axons were considered to be DAT-positive when one or more silver-intensified gold (SIG) particles could be detected along its plasma membrane or within its cytosol. Axons were identified to be forming symmetrical or asymmetrical synapses. SIG particles were categorized as either membranous (center of particle was perfectly aligned to plasma membrane) or intracellular (center of particle was within the boundaries of the axon). The size of the axons and SIG particles were assessed by measuring the maximum and minimum diameters. The distance of each SIG particle to the plasma membrane was measured using the center of the SIG particle and the nearest point along the membrane. 897 SIG particles were included in the analysis.

#### Estrogen receptor suppression

AAV2-U6/TO-shEsr1-CMV-TetR-P2A-eGFP-KASH-pA was custom constructed from AAV backbone (AddGene #82706, also see [64]) with the insertion of the shRNA sequence of *Esr1*^65^ and custom prepped by Wz Biosciences, Inc. pJEP11-AAV-U6/TO-sgRNA(Tet2)-CMV-TetR-P2A-eGFP-KASH-pA was a gift from Jonathan Ploski (AddGene plasmid #82705). shEsr1 was also cloned into the lentiviral vector pLL3.7 as previously described^66,67^ (AddGene plasmid #120720). Either AAV-U6TO-shEsr1-CMV-TetR-eGFP (U6/tetO-shEsr1, *N* = 5) or pLL3.7-shEsr1 (lenti-shEsr1, *N* = 3), which use the same shEsr1 sequence^65^, was injected bilaterally into the VTA in 8 rats to knockdown *Esr1* expression. Doxycycline (dox) was used to induce viral expression of U6/tetO-shEsr1, delivered via both ad lib food (Bio-Serv diet) and 10-20 minutes of ad lib dox-containing water (∼ 3 mg/mL Sigma-Aldrich dox powder) such that each rat received 20 mg/kg per day (below toxic levels)^68^. All behavioral sessions after the first day of the dox regimen were included in the analysis for U6/tetO-shEsr1-injected rats and after 21 days of expression for lenti-shEsr1-injected rats and compared to sessions that preceded surgery.

#### Serum osmolality

Osmolality was measured in the same serum samples used to detect 17*β*-estradiol expression. The extracted serum was sent by refrigerated transportation to a commercial laboratory for osmolality determination using a freezing-point method (Cornell University Clinical Pathology Laboratory).

#### Histological verification with protein and RNA staining

Rats were euthanized following completion of behavioral testing for photometry, optogenetics, and shRNA experiments and when they reached adulthood (10 - 11 weeks) for electron microscopy and immunohistochemistry quantification experiments. Rats were first perfused with phosphate buffer saline (PBS) and subsequently with 4% paraformaldehyde for DAT and SERT quantification, and 10% formalin for histological verification for optogenetics, photometry, and shEsr1 experiments. Brains were post-fixed with 4% paraformaldehyde for one week at 4°C for DAT and SERT quantification, with 10% formalin for two days for verification of surgical coordinates, and with 10% formalin for one day for shEsr1 validation. 50 *µ*m sections of extracted brain tissue were sliced on a Leica VT1000 vibratome.

Sections were treated with 1% hydrogen peroxide in 0.01 M PBS for 30 min, washed with 0.01 M PBS, and blocked in 0.01 M PBS containing 0.05% sodium azide and 1% bovine serum albumin (PBS-BSA-Azide) for 30 minutes before being treated with primary antibody overnight at room temperature (RT). Primary antibody was removed with PBS rinses and then sections were treated with secondary antibodies for 1 hour away from light. Primary and secondary antibodies were made in PBS-BSA-Azide and primary antibodies additionally contained 0.2% triton-x. To verify virus expression and implant location, endogenous fluorescent markers were amplified with rabbit anti-GFP (Thermo Fisher Scientific A11122, 1:2,000, lot #2083201) and imaged with an Olympus VS120 Virtual Slide Microscope. For light microscopy experiments, DAT was labeled with 1:500 Sigma-Aldrich D6944 lot #0000124874, SERT was labeled with 1:2,000 Chemicon AB1594P lot #18020403, TH was labeled with 1:400 Millipore MAB318 lot #3990619, and ER*α* was labeled with 1:250 Thermo Fisher Scientific PA1-309 lot #YG378288. To increase the immunoreactivity of ER*α* in the VTA after fixation, antigen retrieval was conducted with citrate buffer. The secondary antibodies used were as follows (all 1:200 and from Thermo Fisher Scientific): anti-rabbit AlexaFluor 488 IgG A11008, lot #2147635) for DAT, SERT, and GFP, goat anti-mouse AlexaFluor 555 for TH (A21424, lot #2390715), and goat anti-rabbit AlexaFluor 594 (A11037, lot #2160431) for ER*α*. All slices were mounted on glass slides using Prolong Gold (Vector Laboratories).

For verification of *Esr1* knockdown by U6/tetO-shEsr1 and lenti-shEsr1 viruses, RNA fluorescence in situ hybridization (FISH) was performed using the ACD Bio RNAscope Multiplex Fluorescent v2 assay. Brains were cryoprotected in a solution of 10% sucrose/PBS and stored at 4°C until the brain sank. Cryoprotection was repeated with 20% and 30% sucrose/PBS solutions. The brains were embedded in optimal cutting temperature (OCT) solution and flash frozen with 2-methylbutane cooled to -80°C. For tissue sectioning, a cryostat was used to collect 10 *µ*m sections stamped onto superfrost plus slides. The sections were air dried at -20°C for one hour and stored at -80°C. Sections were stained with target probes (EGFP-O4-C1 (ACD Bio 538851, lot #24117A) to label GFP and Rn-Esr1-C2 (ACD Bio 317151-C2, lot #24143D) diluted to 1:50 to label *Esr1*. RNAscope Multiplex Fluorescent Reagent Kit v2 (ACD Bio 323270) was used for the FISH assay. Control tissue was labeled with 3-Plex Positive Control Probe (ACD Bio 320891, lot#24121A) and 3-Plex Negative Control Probe (ACD Bio 320871, lot #2021233) as internal controls. DAPI was used as a counter-stain.

#### Quantification of fluorescence

Photomicrographs were taken at 20x and 63x with a Leica DM6 CS Confocal using standardized parameters developed to maximize signal-to-noise ratios. Z-stacks were taken for *Esr1* quantification at a step size of 0.3 *µ*m.

Photomicrographs were analyzed in ImageJ to quantify protein expression. A mask was generated to identify the location of fluorescing pixels. The total area of the mask (*µ*m^2^) was used to determine expression level of DAT and SERT.

FISH-quant was used for *Esr1* quantification^69^. Detection settings were set based on filtering of background noise and accurate identification of spots on internal control sections, and standardized across all sections. The total number of spots in each image of a z-stack was used for analysis.

### Statistical analyses

#### General

Effect sizes (Cohen’s *d*) for effects across the estrous cycle, which had matching sample sizes across groups, were calculated as:

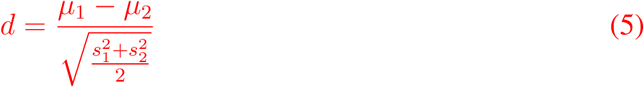

For independent samples, Cohen’s *d* was calculated as:

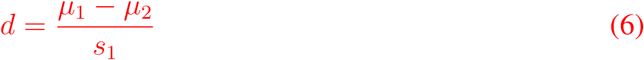

#### Trial initiation times

Trial initiation time was defined as the duration of time in between the termination of one trial (violation, opt out, or the last lick from the rewarded port) and the subsequent center poke that began the next trial. We excluded outlier trial initiation times that were above the 98*^th^* percentile of the rat’s cumulative trial initiation time distribution pooled over sessions, except when less than three sessions per stage were used, then the cut off was 99*^th^* percentile. Sensitivity of trial initiation times to blocks was defined as change in initiation time (mean in low - mean in high blocks). Where noted, the effect of satiety over the session on trial initiation times was accounted for by subtracting the regression of mean trial initiation time as a function of trial number.

#### Trial history effects

To assess trial initiation time sensitivity to previous offers, we regressed trial initiation time against previous rewards. We focused only on mixed blocks. Reward offers from a different block (e.g., a previous high block) were given NaN values. Then, we regressed against the previous nine reward offers, not including the current trial, along with a constant offset. We set the reward for violation, catch, and opt-out trials to zero, since rats do not receive a reward on these trials. We used Matlab’s builtin regress function to perform the regression. With the coefficients, we found the first non-significant coefficient (coefficient whose 95% confidence interval contained zero), and set that coefficient and all following coefficients to zero, as these coefficients cannot meaningfully be interpreted as being different from zero. To determine the relationship between the estrous cycle and the effect of trial history on initiation times, we created a multiple linear regression model that included cycle stage in addition to reward history. When only two stages were considered, diestrus was set to 1 and proestrus was set to 2. When all four stages were considered, diestrus was set to 1, metestrus was set to 2, estrus was set to 3, and proestrus was set to 4 to parallel the cycle of 17*β*-estradiol and the distance from its peak.

#### Wait times

Wait times on catch trials were included in analysis. Wait times *>* 3× the standard deviation above the mean were considered outliers and excluded.

#### Photometry

Trials that followed violations were excluded from the data presented in Fig. 2.

#### Optogenetics

Only control sessions that occurred before the first day of optogenetic stimulation were included in the analysis because optogenetic activation of dopamine terminals had effects on behavior that lasted beyond the day of stimulation. Initiation times were normalized to high blocks in control sessions for both optogenetic and sham groups.

#### Proteomics

Proteins were determined to be significantly differentially expressed with a t-test and a Benjamini-Hochberg false discovery rate (FDR) of 5% to correct for multiple comparisons. Significantly differentially expressed protein signatures identified with mass spectrometry were subjected to functional annotation. STRING database v12 was used to assess enrichment of gene ontology terms for rattus norvegus using a hypergeometric test and all identified proteins as background. According to Geneontology.org, 246 genes and their proteins are associated with dopamine. 92 of these proteins were identified by mass spectrometry and three that were significantly decreased in expression are related to dopamine uptake and transport (Slc6a3, Slc6a4, and Smpd3).

#### Serum osmolality

Osmolality measurements *>* 2.5× the standard deviation above the mean were considered outliers and excluded. 17*β*-estradiol expression for one rat in estrus was excluded when correlated with osmolality due to it being an outlier (*<* 1 standard deviation below the mean of all samples).

## Data Availability

The data generated in this study will be deposited in a Zenodo database upon publication.

## Code Availability

Code used to acquire behavioral data, analyze all data, and generate figures will be available at https://github.com/constantinoplelab/published/tree/main upon publication.

## Acknowledgments

We thank members of the Constantinople lab for feedback. We thank research technicians in the Constantinople lab for animal training. We thank Claudia Farb for her assistance with histology and electron microscopy and NYU Langone’s Proteomics Laboratory (SCR 017926) for the mass spectrometry. We thank Ina Baek Bando and Gina Canino for their assistance with histology processing, imaging, and quantifying.

## Funding

This work was supported by a K99/R00 Pathway to Independence Award (R00MH11-1926), a Klingenstein-Simons Fellowship in Neuroscience, and an NIH Director’s New Innovator Award (DP2MH126376) to C.M.C. C.G. was supported by a grant from the Simons Foundation (855332)), F32MH125448, and 5T32MH019524. The mass spectrometric experiments were supported in part by NYU Langone Health, the Laura and Isaac Perlmutter Cancer Center support grant P30CA016087 from the National Cancer Institute, and the NIH Shared Instrumentation Grant 1S10OD010582-01A1 for the purchase of an Orbitrap Fusion™ Lumos™ Tribrid™ mass spectrometer.

## Author Contributions

C.M.C. and C.E.G. designed the study. C.E.G., A.C.M., D.K., and D.H.L. collected and analyzed data. A.M. developed the behavioral model and contributed to data analysis. T.Y. and D.L. created and shared the U6/tetO-shEsr1 virus. A.L. created and shared the lenti-shEsr1 virus. C.A. supervised the electron microscopy. C.E.G prepared the figures. C.E.G. and C.M.C. wrote the manuscript. C.M.C supervised the project.

## Notes

### Competing Interest Statement

The authors have declared no competing interest.

### Summary of Updates

Figures 1, 2, 3, and 6 revised. Supplemental Figures 1, 2, 4, 6, and 10 revised. Supplemental Figures 3, 5, and 7 added. Discussion and Methods expanded.

## References

1. McEwen, B. S. & Milner, T. A. Understanding the broad influence of sex hormones and sex differences in the brain. Journal of neuroscience research 95, 24–39 (2017).

2. Handy, A. B., Greenfield, S. F., Yonkers, K. A. & Payne, L. A. Psychiatric symptoms across the menstrual cycle in adult women: a comprehensive review. Harvard review of psychiatry 30, 100 (2022).

3. Riecher-Rössler, A., Butler, S. & Kulkarni, J. Sex and gender differences in schizophrenic psychoses—a critical review. Archives of women’s mental health 21, 627–648 (2018).

4. Burt, V. K., Altshuler, L. L. & Rasgon, N. Depressive symptoms in the perimenopause: prevalence, assessment, and guidelines for treatment. Harvard Review of Psychiatry 6, 121–132 (1998).

5. Di Paolo, T. Modulation of brain dopamine transmission by sex steroids. Reviews in the Neurosciences 5, 27–42 (1994).

6. Becker, J. B. Gender differences in dopaminergic function in striatum and nucleus accumbens. Pharmacology Biochemistry and behavior 64, 803–812 (1999).

7. Schultz, W., Dayan, P. & Montague, P. R. A neural substrate of prediction and reward. Science 275, 1593–1599 (1997).

8. Waelti, P., Dickinson, A. & Schultz, W. Dopamine responses comply with basic assumptions of formal learning theory. Nature 412, 43–48 (2001).

9. Bayer, H. M. & Glimcher, P. W. Midbrain dopamine neurons encode a quantitative reward prediction error signal. Neuron 47, 129–141 (2005).

10. Cohen, J. Y., Haesler, S., Vong, L., Lowell, B. B. & Uchida, N. Neuron-type-specific signals for reward and punishment in the ventral tegmental area. nature 482, 85–88 (2012).

11. Day, J. J., Roitman, M. F., Wightman, R. M. & Carelli, R. M. Associative learning mediates dynamic shifts in dopamine signaling in the nucleus accumbens. Nature neuroscience 10, 1020–1028 (2007).

12. Kim, H. R. et al. A unified framework for dopamine signals across timescales. Cell 183, 1600–1616 (2020).

13. Steinberg, E. E. et al. A causal link between prediction errors, dopamine neurons and learning. Nature neuroscience 16, 966–973 (2013).

14. Parker, N. F. et al. Reward and choice encoding in terminals of midbrain dopamine neurons depends on striatal target. Nature neuroscience 19, 845–854 (2016).

15. Sharpe, M. J. et al. Dopamine transients are sufficient and necessary for acquisition of model-based associations. Nature Neuroscience 20, 735–742 (2017).

16. Hamid, A. A. et al. Mesolimbic dopamine signals the value of work. Nature neuroscience 19, 117–126 (2016).

17. Marino, M., Galluzzo, P. & Ascenzi, P. Estrogen signaling multiple pathways to impact gene transcription. Current genomics 7, 497–508 (2006).

18. Vasudevan, N. & Pfaff, D. W. Non-genomic actions of estrogens and their interaction with genomic actions in the brain. Frontiers in neuroendocrinology 29, 238–257 (2008).

19. Yoest, K. E., Quigley, J. A. & Becker, J. B. Rapid effects of ovarian hormones in dorsal striatum and nucleus accumbens. Hormones and behavior 104, 119–129 (2018).

20. Cox, J. & Witten, I. B. Striatal circuits for reward learning and decision-making. Nature Reviews Neuroscience 20, 482–494 (2019).

21. Floresco, S. B. The nucleus accumbens: an interface between cognition, emotion, and action. Annual review of psychology 66, 25–52 (2015).

22. Sutton, R. S. & Barto, A. G. Reinforcement learning: An introduction (MIT press, 2018).

23. Joel, D., Niv, Y. & Ruppin, E. Actor–critic models of the basal ganglia: New anatomical and computational perspectives. Neural networks 15, 535–547 (2002).

24. Chen, R. & Goldberg, J. H. Actor-critic reinforcement learning in the songbird. Current opinion in neurobiology 65, 1–9 (2020).

25. Niv, Y., Daw, N. D., Joel, D. & Dayan, P. Tonic dopamine: opportunity costs and the control of response vigor. Psychopharmacology 191, 507–520 (2007).

26. Xu-Wilson, M., Zee, D. S. & Shadmehr, R. The intrinsic value of visual information affects saccade velocities. Experimental Brain Research 196, 475–481 (2009).

27. Wang, A. Y., Miura, K. & Uchida, N. The dorsomedial striatum encodes net expected return, critical for energizing performance vigor. Nature neuroscience 16, 639–647 (2013).

28. Mah, A., Schiereck, S. S., Bossio, V. & Constantinople, C. M. Distinct value computations support rapid sequential decisions. Nature Communications 14, 7573 (2023).

29. Niv, Y., Joel, D. & Dayan, P. A normative perspective on motivation. Trends in cognitive sciences 10, 375–381 (2006).

30. Starkweather, C. K., Babayan, B. M., Uchida, N. & Gershman, S. J. Dopamine reward prediction errors reflect hidden-state inference across time. Nature neuroscience 20, 581– 589 (2017).

31. Mah, A., Golden, C. & Constantinople, C. Mesolimbic dopamine encodes reward prediction errors independent of learning rates. bioRxiv (2024).

32. Witten, I. B. et al. Recombinase-driver rat lines: tools, techniques, and optogenetic application to dopamine-mediated reinforcement. Neuron 72, 721–733 (2011).

33. Millard, S. J. et al. Cognitive representations of intracranial self-stimulation of midbrain dopamine neurons depend on stimulation frequency. Nature Neuroscience, 1–7 (2024).

34. Larsen, M. B. et al. Dopamine transport by the serotonin transporter: a mechanistically distinct mode of substrate translocation. Journal of Neuroscience 31, 6605–6615 (2011).

35. Kim, S. K. et al. Neutral sphingomyelinase 2 induces dopamine uptake through regulation of intracellular calcium. Cellular signalling 22, 865–870 (2010).

36. Watson, C. S. et al. Estradiol effects on the dopamine transporter–protein levels, subcellular location, and function. Journal of molecular signaling 1, 1–14 (2006).

37. Fleming, W., Jewell, S., Engelhard, B., Witten, D. M. & Witten, I. B. Inferring spikes from calcium imaging in dopamine neurons. PloS one 16, e0252345 (2021).

38. Atcherley, C. W., Wood, K. M., Parent, K. L., Hashemi, P. & Heien, M. L. The coaction of tonic and phasic dopamine dynamics. Chemical Communications 51, 2235–2238 (2015).

39. Zhang, L., Doyon, W. M., Clark, J. J., Phillips, P. E. & Dani, J. A. Controls of tonic and phasic dopamine transmission in the dorsal and ventral striatum. Molecular pharmacology 76, 396–404 (2009).

40. Cox, J. et al. A neural substrate of sex-dependent modulation of motivation. Nature Neuroscience, 1–11 (2023).

41. Melikian, H. E. Neurotransmitter transporter trafficking: endocytosis, recycling, and regulation. Pharmacology & therapeutics 104, 17–27 (2004).

42. Chen, R., Furman, C. A. & Gnegy, M. E. Dopamine transporter trafficking: rapid response on demand. Future neurology 5, 123–134 (2010).

43. Miranda, M., Wu, C. C., Sorkina, T., Korstjens, D. R. & Sorkin, A. Enhanced ubiquitylation and accelerated degradation of the dopamine transporter mediated by protein kinase C. Journal of Biological Chemistry 280, 35617–35624 (2005).

44. Vicentic, A., Battaglia, G., Carroll, F. I. & Kuhar, M. J. Serotonin transporter production and degradation rates: studies with RTI-76. Brain research 841, 1–10 (1999).

45. Calipari, E. S. et al. Dopaminergic dynamics underlying sex-specific cocaine reward. Nature communications 8, 13877 (2017).

46. Zhang, D., Yang, S., Yang, C., Jin, G. & Zhen, X. Estrogen regulates responses of dopamine neurons in the ventral tegmental area to cocaine. Psychopharmacology 199, 625–635 (2008).

47. Shughrue, P. J., Lane, M. V. & Merchenthaler, I. Comparative distribution of estrogen receptor-*α* and-*β* mRNA in the rat central nervous system. Journal of Comparative Neurology 388, 507–525 (1997).

48. Creutz, L. M. & Kritzer, M. F. Estrogen receptor-*β* immunoreactivity in the midbrain of adult rats: Regional, subregional, and cellular localization in the A10, A9, and A8 dopamine cell groups. Journal of Comparative Neurology 446, 288–300 (2002).

49. McHenry, J. A. et al. Hormonal gain control of a medial preoptic area social reward circuit. Nature neuroscience 20, 449–458 (2017).

50. Bocklisch, C. et al. Cocaine disinhibits dopamine neurons by potentiation of GABA transmission in the ventral tegmental area. Science 341, 1521–1525 (2013).

51. Morales, M. & Margolis, E. B. Ventral tegmental area: cellular heterogeneity, connectivity and behaviour. Nature Reviews Neuroscience 18, 73–85 (2017).

52. Matsuda, W. et al. Single nigrostriatal dopaminergic neurons form widely spread and highly dense axonal arborizations in the neostriatum. Journal of Neuroscience 29, 444– 453 (2009).

53. Zhu, J & Reith, M. Role of the dopamine transporter in the action of psychostimulants, nicotine, and other drugs of abuse. CNS & Neurological Disorders-Drug Targets (Formerly Current Drug Targets-CNS & Neurological Disorders) 7, 393–409 (2008).

54. Sotnikova, T. D., Beaulieu, J.-M., Gainetdinov, R. R. & Caron, M. G. Molecular biology, pharmacology and functional role of the plasma membrane dopamine transporter. CNS & Neurological Disorders-Drug Targets (Formerly Current Drug Targets-CNS & Neurological Disorders) 5, 45–56 (2006).

55. Redish, A. D. Addiction as a computational process gone awry. Science 306, 1944–1947 (2004).

56. Montague, P. R., Dolan, R. J., Friston, K. J. & Dayan, P. Computational psychiatry. Trends in cognitive sciences 16, 72–80 (2012).

57. Radulescu, A. & Niv, Y. State representation in mental illness. Current opinion in neurobiology 55, 160–166 (2019).

58. Kastner, P et al. Two distinct estrogen-regulated promoters generate transcripts encoding the two functionally different human progesterone receptor forms A and B. The EMBO journal 9, 1603–1614 (1990).

59. Marcondes, F., Bianchi, F. & Tanno, A. Determination of the estrous cycle phases of rats: some helpful considerations. Brazilian journal of biology 62, 609–614 (2002).

60. Haisenleder, D. J., Schoenfelder, A. H., Marcinko, E. S., Geddis, L. M. & Marshall, J. C. Estimation of estradiol in mouse serum samples: evaluation of commercial estradiol immunoassays. Endocrinology 152, 4443–4447 (2011).

61. Creamer, M. S., Chen, K. S., Leifer, A. M. & Pillow, J. W. Correcting motion induced fluorescence artifacts in two-channel neural imaging. PLoS computational biology 18, e1010421 (2022).

62. Santiago, A. N., Lim, K. Y., Opendak, M., Sullivan, R. M. & Aoki, C. Early life trauma increases threat response of peri-weaning rats, reduction of axo-somatic synapses formed by parvalbumin cells and perineuronal net in the basolateral nucleus of amygdala. Journal of Comparative Neurology 526, 2647–2664 (2018).

63. Chen, Y.-W., Wable, G. S., Chowdhury, T. G. & Aoki, C. Enlargement of axo-somatic contacts formed by GAD-immunoreactive axon terminals onto layer V pyramidal neurons in the medial prefrontal cortex of adolescent female mice is associated with suppression of food restriction-evoked hyperactivity and resilience to activity-based anorexia. Cerebral cortex 26, 2574–2589 (2016).

64. De Solis, C. A., Ho, A., Holehonnur, R. & Ploski, J. E. The development of a viral mediated CRISPR/Cas9 system with doxycycline dependent gRNA expression for inducible in vitro and in vivo genome editing. Frontiers in molecular neuroscience 9, 70 (2016).

65. Musatov, S., Chen, W., Pfaff, D. W., Kaplitt, M. G. & Ogawa, S. RNAi-mediated silencing of estrogen receptor *α* in the ventromedial nucleus of hypothalamus abolishes female sexual behaviors. Proceedings of the National Academy of Sciences 103, 10456–10460 (2006).

66. Lasek, A., Janak, P., He, L, Whistler, J. & Heberlein, U. Downregulation of mu opioid receptor by RNA interference in the ventral tegmental area reduces ethanol consumption in mice. *Genes*, Brain and Behavior 6, 728–735 (2007).

67. Satta, R., Certa, B., He, D. & Lasek, A. W. Estrogen receptor *β* in the nucleus accumbens regulates the rewarding properties of cocaine in female mice. International Journal of Neuropsychopharmacology 21, 382–392 (2018).

68. El-Neweshy, M. S. Experimental doxycycline overdose in rats causes cardiomyopathy. International Journal of Experimental Pathology 94, 109–114 (2013).

69. Mueller, F. et al. FISH-quant: automatic counting of transcripts in 3D FISH images. Nature methods 10, 277–278 (2013).

