## Supplemental Figures for "Estrogenic control of reward prediction errors and reinforcement learning"

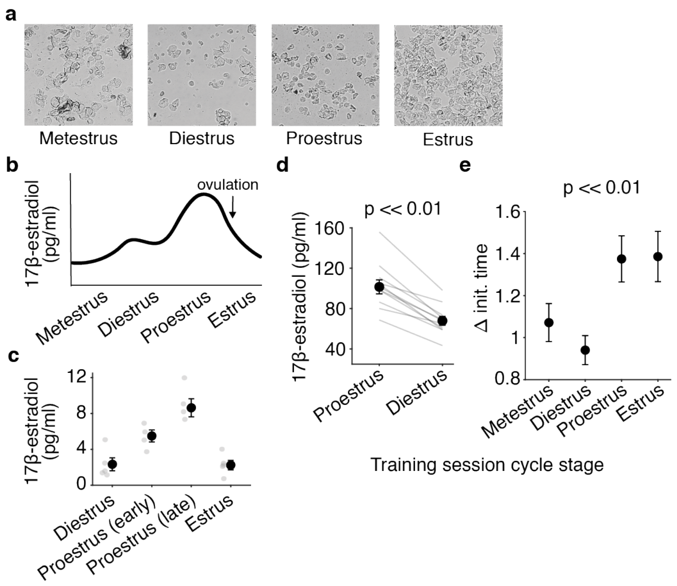


Extended Data 1 : **Trial initiation times are most modulated by the value of the state around the time of ovulation. a.** Example images of vaginal cytology by estrous stage. **b-c.** Expected **(b)** and observed **(c)** median $17\beta$-estradiol, measured with an ELISA from serum. **d.** Serum $17\beta$-estradiol is significantly higher in proestrus than diestrus during training sessions, Wilcoxon signed rank test $p = 9.77 \times 10^{-4}$, $d = 1.79$. **e.** Median sensitivity of detrended trial initiation times to blocks for each estrous stage, Kruskal–Wallis test $p = 5.00 \times 10^{-5}$ for group effect, two-sided Wilcoxon signed-rank tests for post-hoc analyses: $p = 0.002$ and $d = 0.17$ for proestrus vs. metestrus, $p = 2.67 \times 10^{-8}$ and $d = 0.41$ for proestrus vs. diestrus, $p = 0.09$ and $d = 0.17$ for proestrus vs. estrus, $p = 9.06 \times 10^{-11}$ and $d = 0.32$ for estrus vs. metestrus, $p = 2.67 \times 10^{-15}$ and $d = 0.53$ for estrus vs. diestrus, and $p = 9.34 \times 10^{-4}$ and $d = 0.20$ for metestrus vs diestrus). All circles are medians and error bars are median $\pm$ SEM.

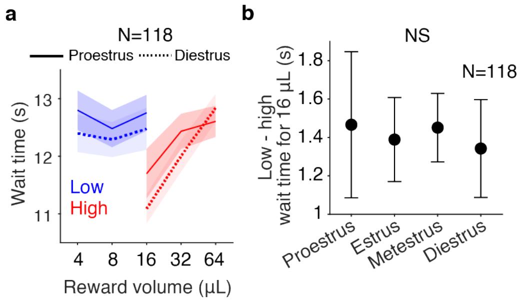


Extended Data 2 : **Wait time adaptation to reward context does not change across the cycle. a.** Mean wait time on catch trials by reward in each block (blue = low, red = high) averaged across rats. Error bars are mean ± SEM. **b.** Median adaptation of wait times to reward blocks (low - high block) does not change across the cycle, Kruskal–Wallis test $p = 0.69$ for group effect and two-sided Wilcoxon signed-rank test $p = 0.771$ and $d = 0.06$ for proestrus vs. diestrus. Error bars are median ± SEM.

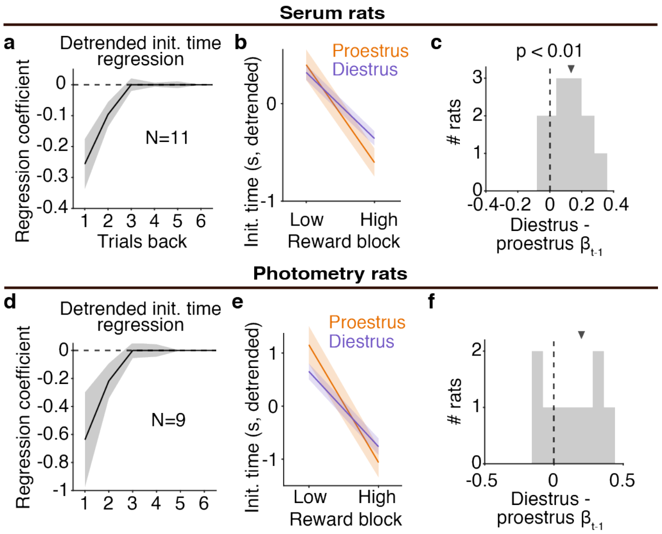


Extended Data 3 : **Population-level effects of the estrous cycle hold in rats used in serum 17$\beta$-estradiol and photometry experiments. a-c.** Rats used in serum 17$\beta$-estradiol experiments have (a) median regression coefficients of detrended trial initiation times plotted against rewards on previous trials during mixed blocks that decline as an exponential function of time, (b) enhanced block sensitivity in proestrus, and (c) greater regression coefficients for the previous reward in proestrus because the distribution is significantly different from zero using a Wilcoxon signed-rank test $p = 4.88 \times 10^{-3}$. **d-f.** Rats used in photometry experiments have (d) median regression coefficients of detrended trial initiation times plotted against rewards on previous trials during mixed blocks that decline as an exponential function of time, (e) enhanced block sensitivity in proestrus, and (f) greater regression coefficients for the previous reward in proestrus. All arrows are medians and all error bars are median ± SEM.

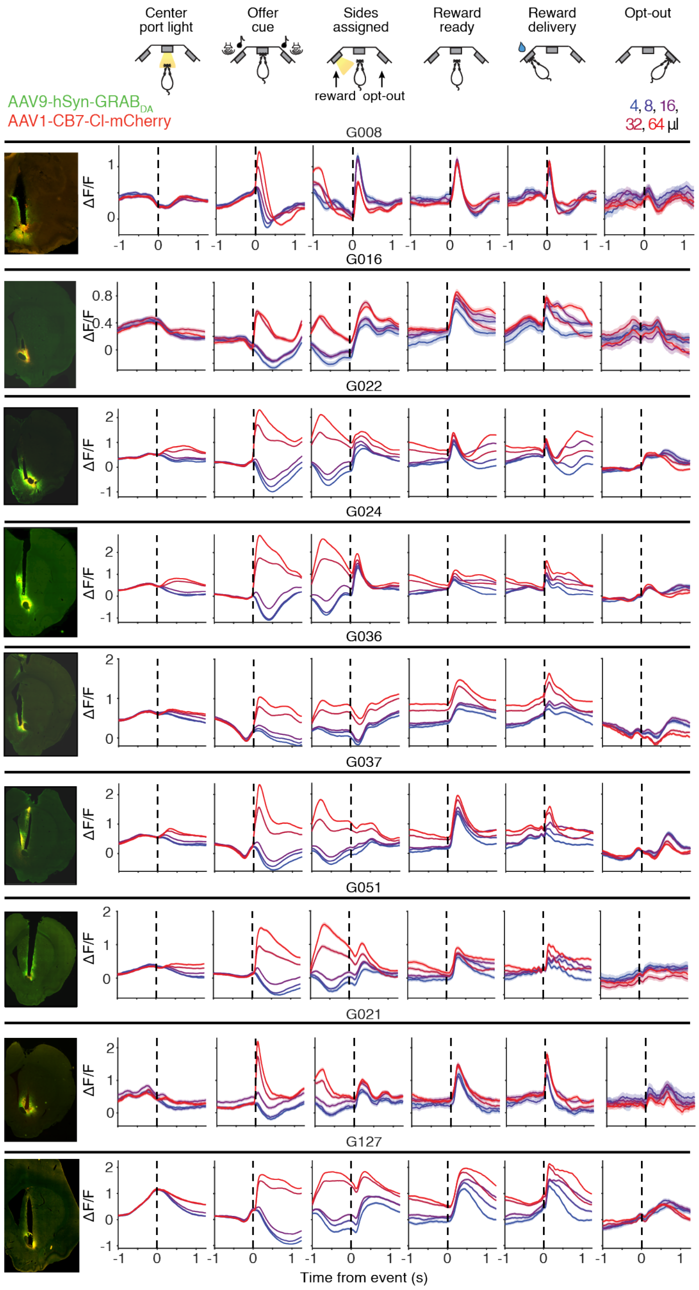


Extended Data 4 : **Histology and task-aligned dopamine across 9 individual rats.** For each rat, histological verification of fiber implant targeting and mCherry (red) and ChR2 (green) expression (left) and task event-aligned dopamine responses, split by reward offer cue volume (right). Data is not baseline-corrected. All error bars are mean ± SEM.

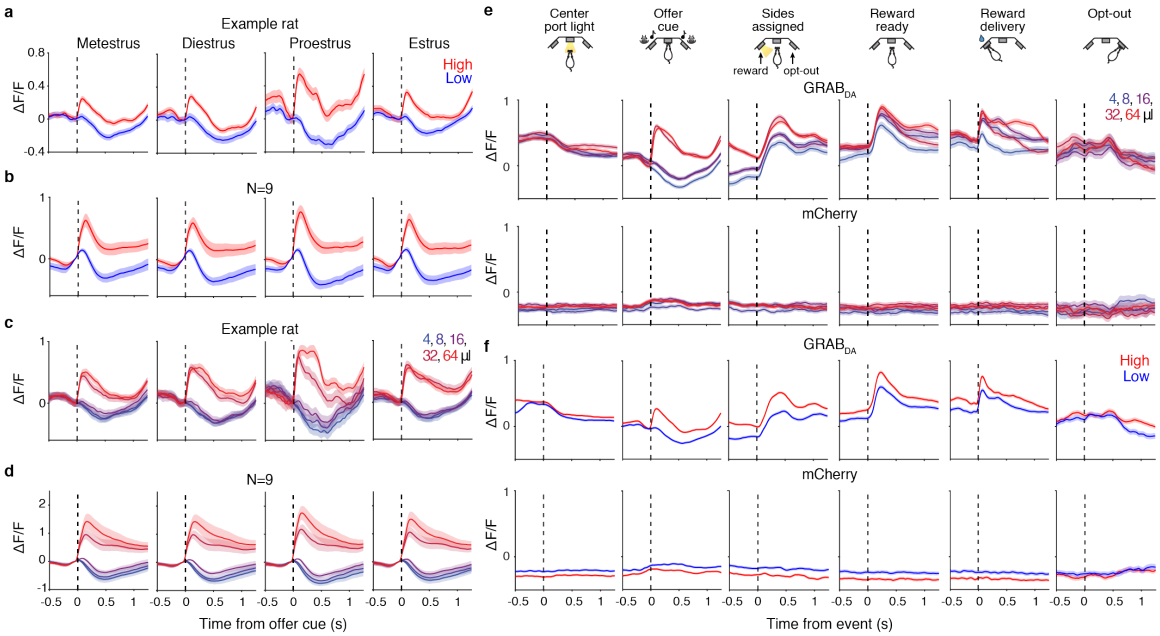


Extended Data 5 : **RPE is most strongly encoded in proestrus compared to the rest of the estrous cycle and not encoded by mCherry.** Baseline-corrected mean dopamine response to reward offer cue split by block for an example rat (**a**) and across the population (**b**), and split by reward volume for an example rat (**c**) and across the population (**d**). **e.** While motion-corrected $GRAB_{DA}$ encodes reward volume, mCherry, used to control for motion artifacts, does not. **f.** Same for block encoding.

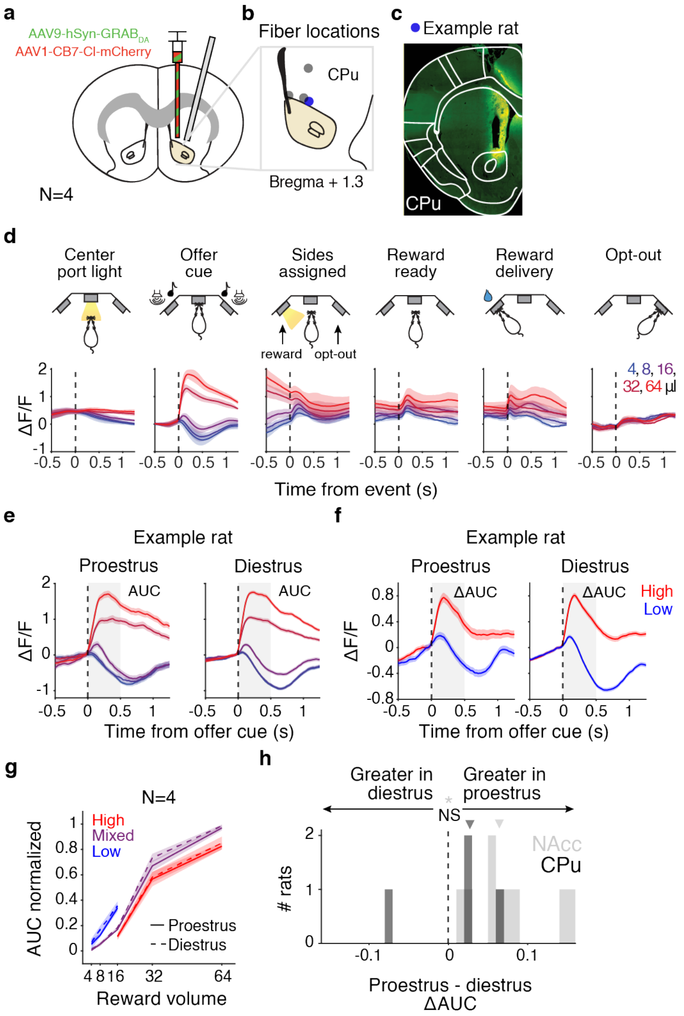


Extended Data 6 : **Dopamine in caudate putamen above NAcc encodes RPE, but is unaffected by estrous. a-b.** Fiber locations for 4 rats. Blue circle in **b** is example depicted in **c**. CPu = caudate putamen. **c.** Example histology of blue circle in **b**. **d.** Task event-aligned dopamine signaling, separated by reward volume. **e-f.** Response to offer cue, separated by reward volume **(e)** and block **(f)** and stage group for example rat, G027. Gray boxes represent the window used to calculate change in area under curve, 0 to 0.5 s (ΔAUC) in **(g)** and **(h)**. Data is baseline-corrected using the 0.05 to 0 s before offer cue. **g.** No significant difference in median population response to each reward volume, separated by reward block, min-max normalized, and baseline-corrected using the 50 ms before offer cue. **h.** Histogram of stage effect (proestrus-diestrus) on ΔAUC during offer cue for all NAcc and CPu rats ($*p = 3.9 \times 10^{-3}$ for NAcc using a two-sided Wilcoxon signed-rank test, $p = 0.875$ for CPu). Triangles are medians. All error bars are mean ± SEM.

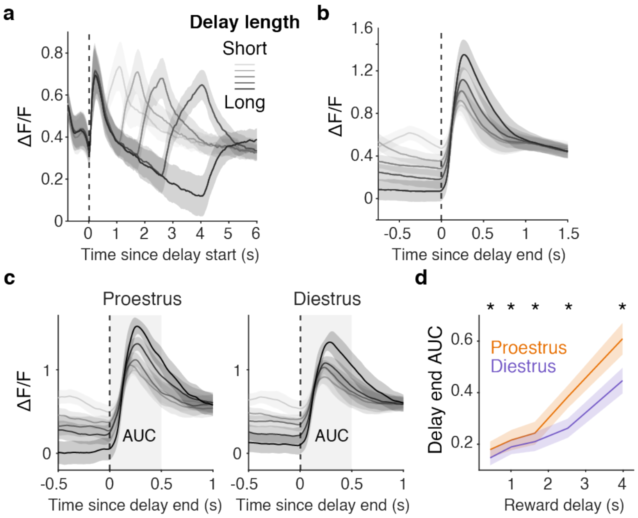


Extended Data 7 : **Delay end RPE is enhanced in proestrus. a.** Mean dopamine response aligned to beginning of delay period for rewarded mixed block trials by reward delay length. **b.** Mean dopamine response aligned to the end of the delay period for rewarded mixed block trials by reward delay length. **c.** Dopamine response aligned to the end of the delay period when reward was cued for mixed block trials by reward delay length and stage. **d.** Median AUC from c is greater in proestrus than diestrus, Wilcoxon signed-rank test. $N = 9$, *, $p < 0.05$. Error bars are ± SEM.

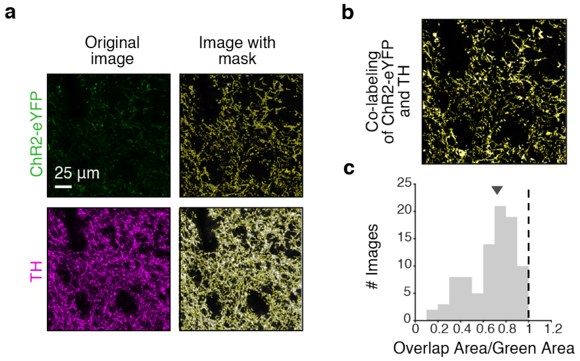


Extended Data 8 : **Expression of ChR2 in TH+ axons terminals in the NAcc. a.** Example images from immunohistochemical quantification of ChR2-eYFP (top) and TH (bottom) co-labeling at 63x with mask (right) identifying fluorescent pixels above a threshold, generated in ImageJ, overlaid. **b.** Image created in ImageJ that depicts pixels that are fluorescent in both ChR2-eYFP and TH images (overlap) with mask. **c.** Metric of co-labeling across all images: area of mask segmentation for ChR2-eYFP image divided by area of mask segmentation for TH image. The arrow indicates the median.

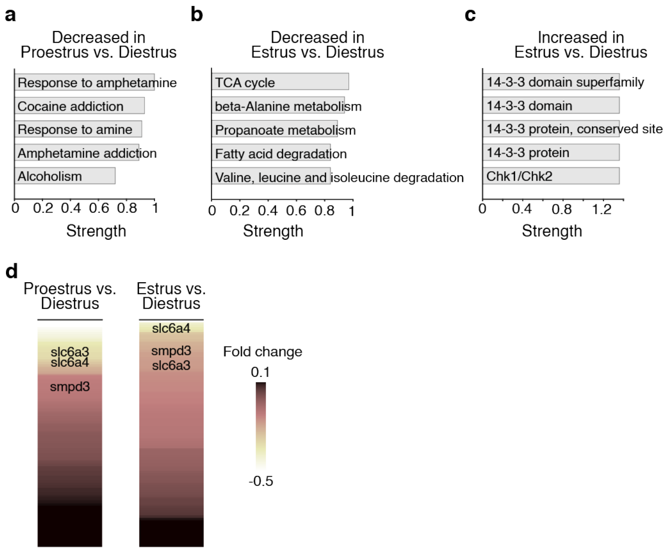


Extended Data 9 : **Proteins related to dopamine reuptake are significantly decreased in proestrus and estrus. a-c.** The strongest gene ontology terms that decreased **(a-b)** and increased **(c)** proteins were significantly enriched for in proestrus **(a)** and estrus **(b-c)** compared to diestrus. Increased proteins in proestrus compared to diestrus were not significantly enriched for any known biological functions. **d.** 92 differentially expressed proteins related to dopamine sorted by fold change, with proteins involved in dopamine reuptake identified.

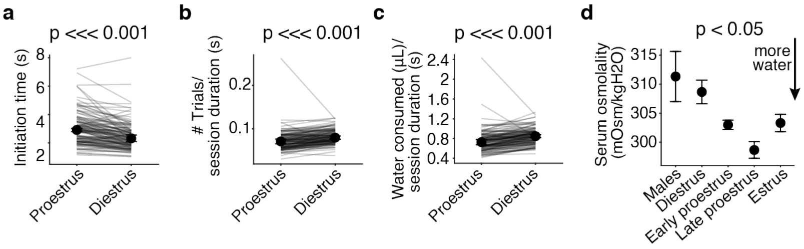


Extended Data 10 : **Systemic expression of 17$\beta$-estradiol reduces thirst. a.** Median initiation time across all trials is significantly higher in proestrus, two-sided Wilcoxon signed rank test $p = 1.60 \times 10^{-14}$ and $d = 0.49$ ($N$ = 118). **b-c.** Median trials performed (**b**) and total water consumed (**c**), controlling for the total amount of time the rat was allotted in the behavioral rig ($N$ = 118). **d.** Median serum osmolality was lower for stages around ovulation (proestrus and estrus, $N$ = 5 males, $N$ = 7 diestrus, $N$ = 4 early proestrus, $N$ = 4 late proestrus, and $N$ = 5 estrus), Kruskal–Wallis test $p = 0.01$ for the group effect and $d = 1.61$ for proestrus vs diestrus.
